# Self-organised pattern formation in the developing neural tube by a temporal relay of BMP signalling

**DOI:** 10.1101/2023.11.15.567070

**Authors:** S Lehr, D B Brückner, M Greunz-Schindler, T Minchington, J Merrin, E Hannezo, A Kicheva

**Author notes:** Equal contribution (S Lehr, D B Brückner).

## Abstract

Developing tissues interpret dynamic changes in morphogen activity to generate cell type diversity. To quantitatively study BMP signalling dynamics in the vertebrate neural tube, we developed a new ES cell differentiation system tailored for growing tissues. Differentiating cells form striking self-organised patterns of dorsal neural tube cell types driven by sequential phases of BMP signalling that are observed both in vitro and in vivo. Data-driven biophysical modelling showed that these dynamics result from coupling fast negative feedback with slow positive regulation of signalling by the specification of an endogenous BMP source. Thus, in contrast to relays that propagate morphogen signalling in space, we uncover a BMP signalling relay that operates in time. This mechanism allows rapid initial concentration-sensitive response that is robustly terminated, thereby regulating balanced sequential cell type generation. Altogether, our study provides an experimental and theoretical framework to understand how signalling dynamics are exploited in developing tissues.

## Introduction

During development, a small number of morphogens are reused in multiple tissues to control pattern formation and tissue growth. While morphogens are commonly viewed as molecules that determine cell fate responses in a concentration-dependent manner, it is becoming increasingly clear that the temporal dynamics of signalling are also relevant for pattern formation ^1–3^. Multiple temporal features, such as duration of signalling or rate of change of signals over time, have been linked to downstream responses ^4–7^. Nevertheless, in many systems how the temporal dynamics of signalling are controlled and linked to cell fate decisions is unclear.

In the developing mouse spinal cord, pattern formation along the dorsal-ventral (DV) axis occurs in response to opposing gradients of BMP and Shh signalling ^8^. In the dorsal spinal cord, the acquisition of distinct neural progenitor identities has been proposed to depend on the BMP levels, signalling duration or ligand type ^9–12^. Yet, it has been challenging to link these dependencies to the dynamics of the endogenous BMP signalling gradient. The spatiotemporal profile of BMP signalling in mouse has been measured in the closed neural tube ^13,14^, but the earlier dynamics and the mechanisms that underlie the establishment of the gradient are poorly understood. BMP ligands are first expressed within the surface ectoderm adjacent to the neural epithelium and later their expression is initiated within the dorsal-most cells of the neural tube, termed the roof plate ^15,16^. BMP ligands produced by both of these sources signal to cells within the developing neural tube.

Besides the changing geometry of the sources of ligand production, the BMP signalling gradient within the neural tube is established in the context of ongoing pattern formation and tissue growth, as well as morphogenetic changes resulting from the specification and emigration of the neural crest (NC). The most dorsal region of the posterior neural tube is at first composed of progenitors that give rise to trunk neural crest. Soon after their specification, neural crest progenitors undergo EMT and migrate out of the neural tube. The remaining progenitors give rise to spatially ordered domains consisting of roof plate (RP), marked by Lmx1a expression, and dorsal neural progenitor subtypes dp1-6 (Fig. 1A) (reviewed in ^17–19^). These observations therefore raise the question how the sequential dynamics of neural crest and roof plate and neural progenitor specification interplays with the dynamics of the BMP signalling gradient.

**Figure 1.**
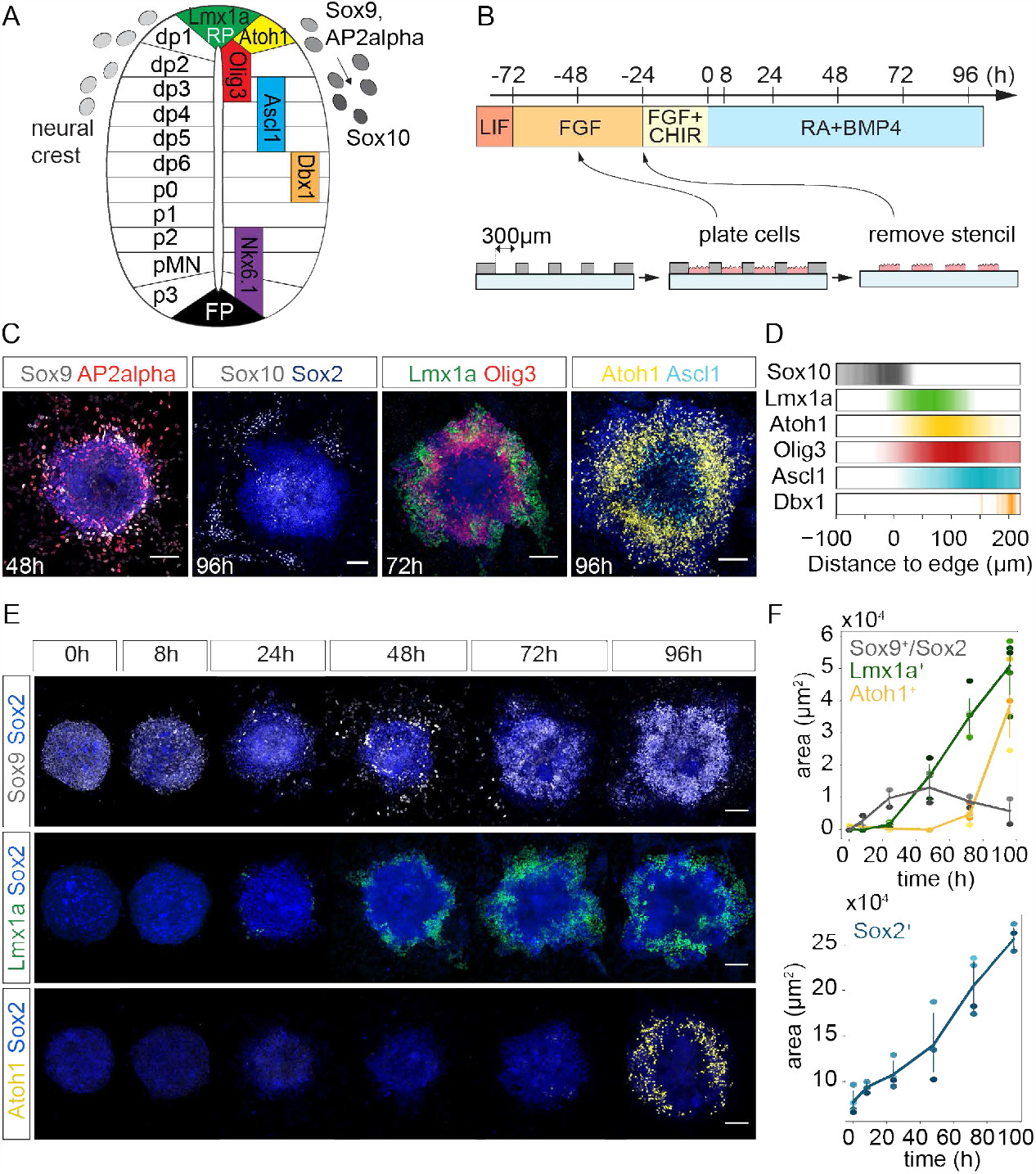
2D stencil differentiation system captures dorsal neural tube patterning. **A**) Cell types and gene expression pattern in the E10.5 neural tube. RP roof plate, FP floor plate. **B)** Differentiation protocol used in this study (top). t=0 is the time when exogenous 0.5 ng/ml BMP4 is added, unless otherwise indicated. Illustration of the stencil method (bottom). **C, E)** Immunostainings against the indicated markers shows self-organised patterns with NC (Sox9, AP2alpha, Sox10) and RP (Lmx1a) at the periphery. Scale bars 100µm. **D)** Quantifications of gene expression profiles from immunostainings at 96h. Colour intensity corresponds to mean FI of n=23-38 colonies except for Dbx1 where n=10. **F)** Mean expression area of the indicated genes quantified from immunostainings. Dots are means of individual experiments, line is mean of all experiments. n=26-79 (Lmx1a), n=8-64 (Atoh1), n=12-22 (Sox9), n=25-53 (Sox2) colonies per gene per time point. Error bars, 95% CI.

Here we establish a novel in vitro system of two-dimensional mouse ES cell differentiation using stencil-based micropatterning to study morphogen-driven pattern formation of the dorsal neural tube. This system, unlike commonly used micropattern techniques ^20–22^, allows tissue growth and cell migration. Our differentiation protocol yields remarkably reproducible self-organised patterns of dorsal neural tube cell types upon exposure to BMP. Strikingly, we found that cells respond to a BMP signalling gradient that is not constant in time, but is activated in two sequential phases. A biophysical model together with perturbation experiments showed that these biphasic signalling dynamics result from a network architecture with interconnected negative and positive feedback loops that work on different time scales. Our analysis shows that the transcription factor Lmx1a is a key mediator that connects the two signalling phases and relays the signal from the initial environmental input to the formation of an endogenous source of BMP production. This relay mechanism combines a rapid concentration-sensitive response that initiates neural crest and roof plate formation with a built-in timer that allows the subsequent reuse of the pathway for patterning of the dorsal neural progenitor subtypes. Our study reveals how the response of cells to a morphogen self-generates complex spatiotemporal morphogen dynamics that robustly encodes cell diversity.

## Results

### In vitro differentiated dorsal neural tube progenitors self-organise following a defined spatiotemporal sequence

Directed differentiation of embryonic stem cells provides a tractable system to investigate how cells interpret signals to generate defined cell identities in a characteristic spatiotemporal order. Cells of the posterior neural tube (Fig. 1A) arise from neuromesodermal progenitors (NMPs) ^23,24^. To generate these cell types in vitro, we therefore based our approach on a 2D monolayer protocol for generating mouse NMPs by transient exposure to the Wnt agonist CHIR99021 ^25^. BMP4 is known to be a dorsalising factor in the developing mouse neural tube ^15^, hence we reasoned that exposure of NMPs to BMP4 in parallel with retinoic acid (RA), which promotes neural identity, will result in the generation of dorsal neural tube progenitors (Fig. 1B, top). Remarkably, we found that exposure of the NMPs to 0.5 ng/ml BMP4 resulted in the formation not of a single cell type, but rather of all dorsal neural tube progenitor types in their correct spatial order, with neural crest forming at the periphery of colonies and dorsal neural progenitors in the centre (Fig. 1C, D, S1A). RNA sequencing analysis of differentiated colonies at 24 and 48h revealed that they express genes characteristic of neural crest, roof plate, as well as dorsal neural progenitor domains dp1-6 (Fig. S1B). The expression of Hox paralogues 3 to 9 indicated that differentiated cells had axial identities corresponding to posterior hindbrain, brachial and thoracic spinal cord levels (Fig. S1C).

To quantitatively characterise these self-organised patterns, we sought to optimise the reproducibility of pattern formation in our protocol. Combining ES cell differentiation with geometric constraints, such as culture on micropatterned surfaces, is known to reduce heterogeneity and improve the reproducibility of patterning ^26^. However, culture on micropatterned surfaces restricts colony growth to a predefined size, which in our case resulted in the formation of often irregular three-dimensional structures with clusters of Erbb3 marking neural crest cells at the periphery (Fig. S1D). To circumvent this, we established a protocol to initialise colonies on a defined geometry and subsequently allow neural crest migration and colony expansion as a monolayer (Methods, ^27^). To do this, we seed the cells on microfabricated stencils made of thin silicone sheets with circular thru holes 300μm in diameter. After ∼20h, just prior to the CHIR pulse, stencils are removed (Fig. 1B, bottom). This approach yields (2D) colonies with migrating neural crest at the periphery and a Sox2+ neural progenitor core that robustly increases in size by ∼5 fold over the course of 96h after BMP4 addition (Fig. 1C-F).

To understand how this self-organised pattern forms, we analysed the spatiotemporal profiles of gene expression over the course of 4 days after BMP4 addition by immunofluorescence. At the periphery of colonies, we observed early migratory neural crest cells, marked by AP2alpha and Sox9 expression (Fig. 1C, E). These cells could be observed as early as 8h after BMP4 exposure (Fig. 1E, F). From 24h onwards, Sox10 expressing late migratory neural crest cells were observed (Fig. 1C, D, S1B). RP and genes characteristic of the dorsal neural progenitor domains dp1 to dp6 formed nested concentric rings that were positioned from the periphery to the centre following the same spatial order as in vivo (Fig. 1C-F, S2A, B). qPCR (Fig. S1E) and immunostaining (Fig. 1E, F) analysis showed that Lmx1a expression characteristic of the RP is detectable after 24h, while Atoh1, which is expressed by dp1 progenitors appears first at 72h. The temporal order of formation of these cell types is therefore consistent with their formation in vivo (Fig. S2A, B).

### Biphasic dynamics of BMP signalling underlies self-organised cell fate patterning

Because dorsal neural tube patterning depends on BMP signalling, the self-organisation of cell fate patterns that we observe suggests the formation of a BMP signalling gradient in the cell colonies. To test this, we performed an immunostaining against phosphorylated Smad1/5 (pSmad1/5), a direct readout of BMP signalling, at different time points of differentiation. This analysis revealed that pSmad1/5 has a graded profile of activity by 8h, with the highest levels observed at the periphery of colonies (Fig. 2A-C). Surprisingly, however, we found that pSmad1/5 activation does not persist continuously over time, but instead occurs in two distinct phases. Initially, a pSmad1/5 signalling gradient appears rapidly, within 2h of BMP addition (Fig. 2C). Subsequently, by 24h pSmad1/5 activity declines to background levels. At 48h, pSmad1/5 is upregulated again in a graded manner from the margin reaching maximum levels from 72h onwards (Fig. 2A-C). This observation raises the question how such biphasic dynamics of BMP signalling is regulated and related to cell fate patterning.

**Figure 2.**
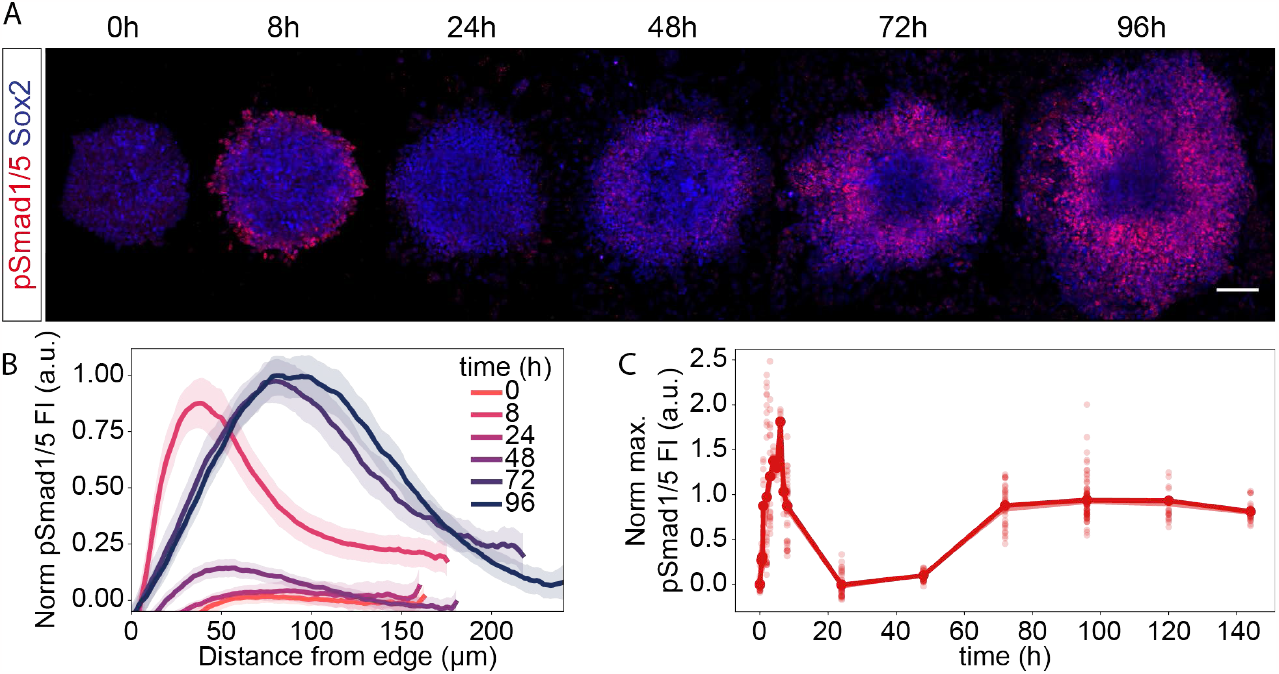
Biphasic temporal dynamics of pSmad1/5 signalling. **A**) pSmad1/5 immunostaining of cells treated with 0.5ng/ml BMP4 and harvested at the indicated times. Scale bar 100µm. **B)** Mean spatial profiles of pSmad1/5 fluorescence intensity (FI) over time. Shaded regions, 95%CI. n=36-51 colonies from 4 experiments. **C)** Maximum pSmad1/5 FI of the spatial profiles were determined for each time point and plotted as a timecourse. Data shows mean and 95%CI, sample sizes per time point: for t=0,8-96h n=43-97 colonies from 5 experiments, for t=0.5-7h n=8-31 colonies from 2 experiments, for t=120, 144h n=30-31 from 2 experiments.

To address this, we first asked whether the formation of polarised pSmad1/5 signalling profiles as well as cell fate patterns in our colonies was dependent on the addition of exogenous BMP4. We found that if no BMP is added to the medium, the formation of organised dorsal pattern did not occur. Instead, in a small fraction of colonies, we observed sporadic non-patterned activation of pSmad1/5 and Lmx1a starting at 48h, but no neural crest (Fig. 3A, B, conditions i and ii).

**Figure 3.**
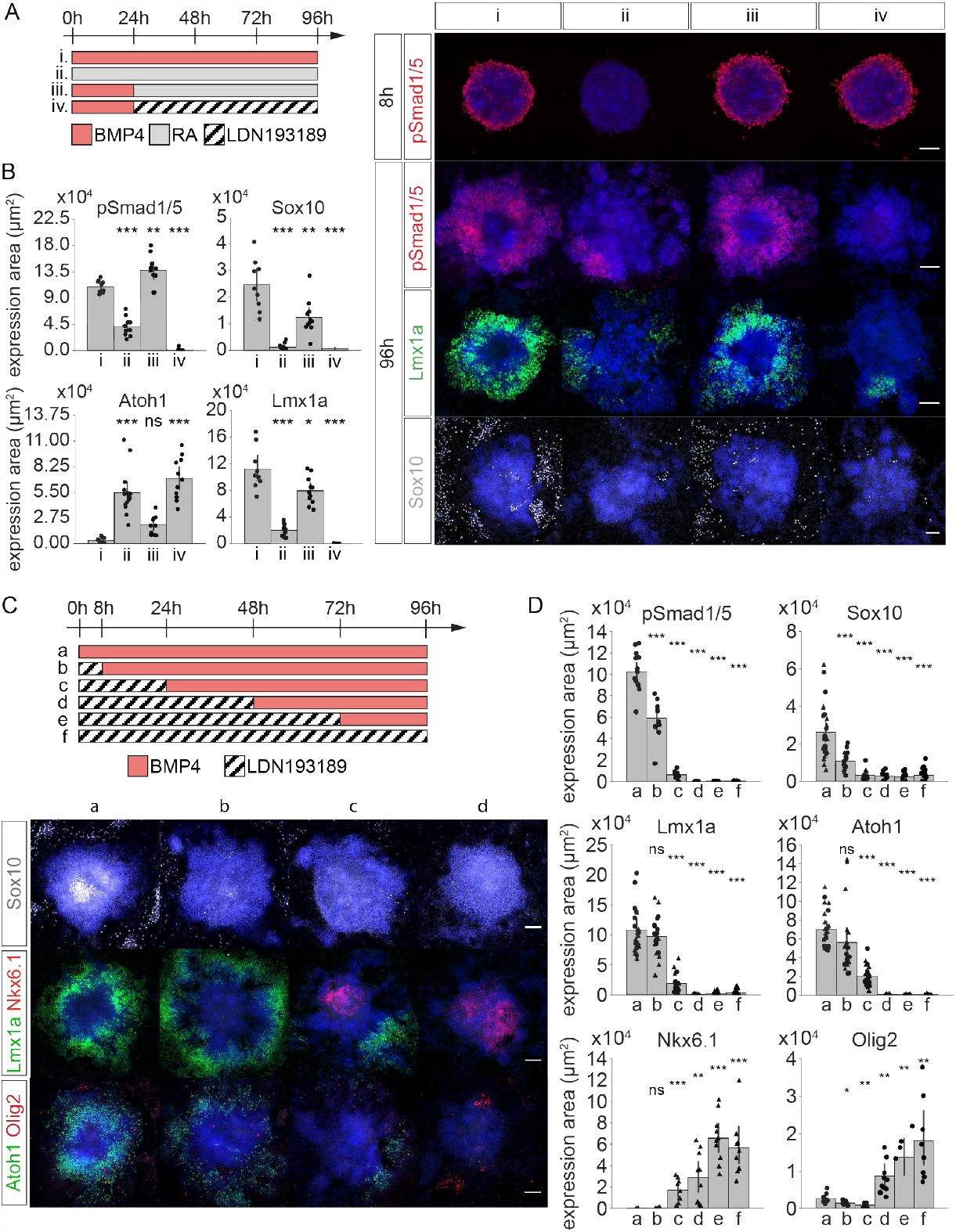
Time window perturbations define the requirements for BMP signalling. **A**) Schematic of differentiation conditions analysed (left). Red indicates RA+0.5ng/ml BMP4, grey only RA, hatched RA + 1μM LDN193189. Immunostainings for the indicated markers and Sox2 (blue) (right). Scale bars 100µm. **B)** Quantification of expression levels (positive area) of the experiment in A. For Sox10 in B, D, the Sox10+, Sox2-negative area was quantified. n=9-15 colonies per condition. Conditions ii-iv were compared to i with t-test: ns p>0.05, *p≤0.05, **p≤0.001, ***p≤0.0001. Data shows mean and 95%CI. **C)** Differentiation conditions, colour code as in A. Representative images (bottom) at 96h. Sox2 in blue. Scale bars 100µm. **D)** Quantification of expression levels (positive area) of the experiment in C. n=13-25 (Sox10), n=18-21 (Lmx1a), n=20-24 (Atoh1), n=8-14 (pSmad1/5), n=9-10 (Nkx6.1), n=8-9 (Olig2). Conditions b-f were compared to a with t-test as in B.

To define critical time windows of BMP signalling for pattern formation, we used the BMP receptor inhibitor LDN193189 to inhibit signalling during defined time intervals. Inhibition of BMP signalling from the beginning of neural differentiation for periods of 24h or longer severely impaired the formation of all dorsal neural tube cell types at 96h, including neural crest, roof plate and dp1 progenitors, marked by Sox10, Lmx1a and Atoh1 expression, respectively (Fig. 3C, D). Instead, an increasing fraction of Sox2 positive neural progenitors adopted ventral identities, marked by Nkx6.1 and Olig2 expression (Fig. 3C, D). BMP inhibition in the first phase (24h) was sufficient to also block subsequent pSmad1/5 signalling in the second phase (Fig. 3D). Thus, the first phase of pSmad1/5 activity is necessary for subsequent pSmad1/5 signalling and for the formation of self-organised cell fate pattern.

To understand whether the first phase of pSmad1/5 is also sufficient for pattern formation, we added BMP to the medium at t=0h for 24h and subsequently replaced it with medium that contained only RA but no BMP. Strikingly, dorsal neural tube pattern formation, assessed by the expression profiles of NC, RP and dp1 markers in this experiment was indistinguishable from pattern formation in the case when BMP exposure was continuous (Fig. 3A, B, conditions i and iii, S3A). Likewise, the profiles of pSmad1/5 activity were similar in the conditions in which BMP4 was added as a 24h pulse or continuously (Fig. 3A, B, S3B). These observations suggested that cell fate patterning and pSmad1/5 activity after the first 24h are independent of the exogenous BMP in the medium, leading us to hypothesise that it depends instead on the endogenous expression of BMP ligands.

To test this hypothesis, we assessed the endogenous expression of BMP ligands over time in our differentiated cells using qPCR. This analysis indicated that the expression of several ligands of the BMP family, including BMP6, BMP7, GDF7 and BMP4 begins at low levels at 24h of differentiation and continuously increases until 96h (Fig. S3C). This is consistent with the possibility that these ligands induce the second phase of pSmad1/5 activity, which is detected from 48h onwards. Consistent with this, inhibition of BMP signalling from 24h onwards using LDN prevents the activation of pSmad1/5 during the second phase (Fig. 3A, B, condition iv). The inhibition of BMP signalling with LDN from 24h or 48h onwards also resulted in a marked decrease in Lmx1a and Sox10 expression (Fig. 3A, B), suggesting that ongoing BMP signalling after the first 24h is necessary for the correct expansion of these cell populations. Together, these results indicate that the first peak of pSmad1/5, which is dependent on exogenous BMP4, is necessary and sufficient to induce a second phase of pSmad1/5 activity by inducing endogenous production of BMP ligands.

### A temporal signalling relay underlies the biphasic dynamics of BMP signalling

To further investigate the mechanisms that underlie the temporal changes in pSmad1/5 levels, we developed a minimal spatiotemporal model of pSmad1/5 signalling (SI). In our model, pSmad1/5 levels depend on the local concentration of BMP ligands with constant sensitivity, except for cells at the edge of the colony, which are more sensitive to BMP (SI). This edge effect is similar to that observed in other micropattern protocols ^22,28,29^ and has been attributed to differential receptor accessibility and mechanical effects. In turn, pSmad1/5 induces the expression of diffusible BMP ligands, based on our observation that endogenous ligand expression is induced upon BMP treatment (Fig. S3C). The observed downregulation of pSmad1/5 at 24h, together with the observation that exogenous signalling is not required to maintain pSmad1/5 signalling levels after 24h (Fig. 3A, B, S3B), implies that a negative regulator is also relevant to the dynamics. Inspection of the RNA sequencing dataset indicated that a number of BMP inhibitors are upregulated in differentiating colonies in response to BMP treatment as early as 8h (Fig. S4A). This is consistent with previous findings that inhibitors of BMP signalling activity are also often targets of the pathway ^30,31^. We therefore also included in the model a generic inhibitor of BMP, which is produced as a function of pSmad1/5 activity and inhibits BMP-mediated pSmad1/5 activation (Fig. 4A).

**Figure 4.**
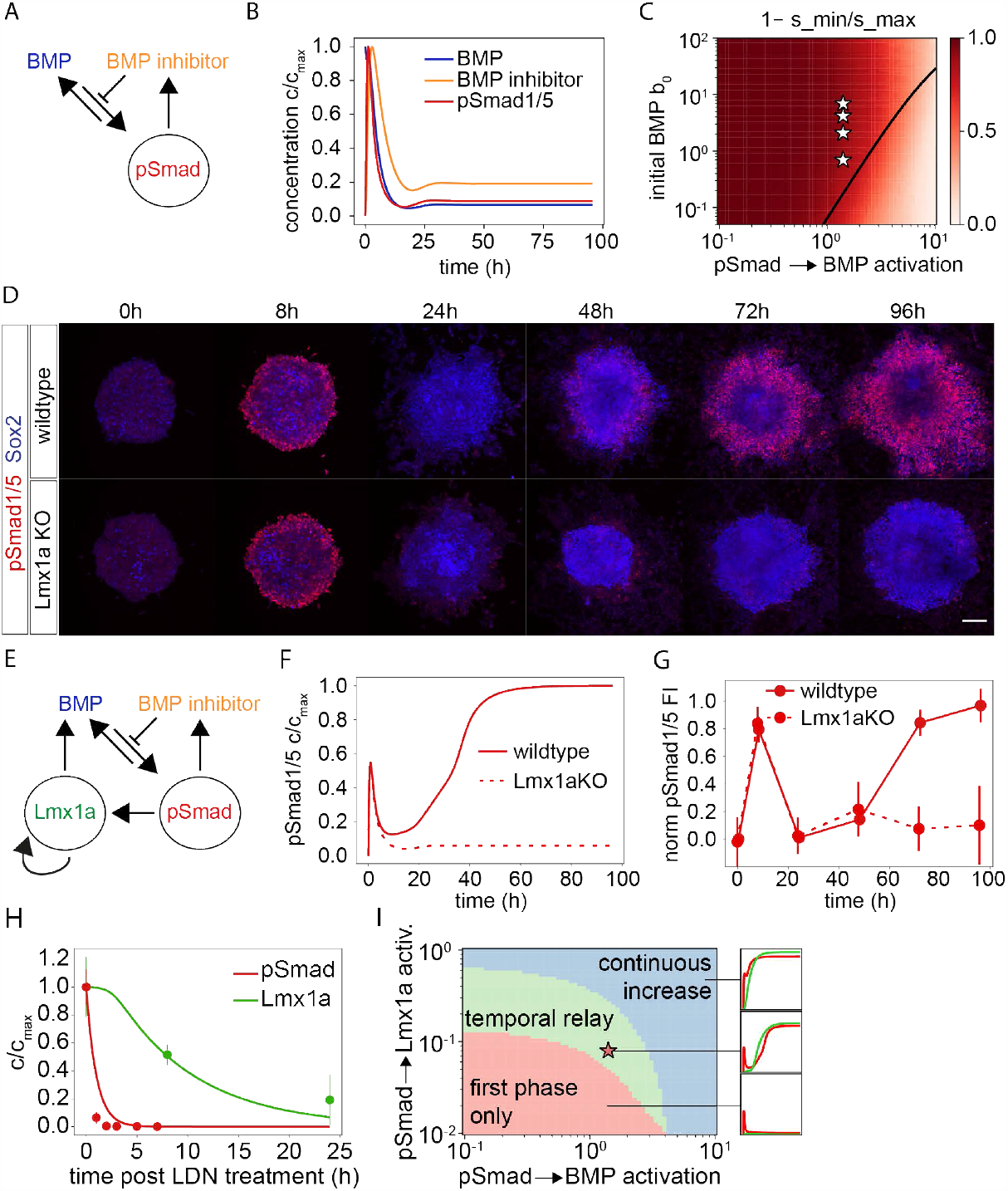
Formulation of a mathematical model of BMP signalling dynamics. **A, B**) Minimal interaction network (A) that captures the first phase signalling dynamics of pSmad1/5 (B). **C)** Downregulation strength (colour-coded, defined as 1-s_min/s_max, where s_max is the max pSmad level in the time course and s_min the subsequent minimum) as a function of initial BMP concentration *b*_0_and the activation strength of BMP by pSmad1/5 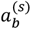. **D)** pSmad1/5 immunostaining in wt and Lmx1aKO cells treated with 0.5ng/ml BMP4. Scale bar 100µm. **E)** Extended interaction network including positive feedback via Lmx1a. **F)** Simulation of the extended model (solid). Lmx1aKO condition (dashed) is simulated by removing all Lmx1a production terms. **G)** Quantification of the experiment in D. Error bars, 95%CI. n=44-67 images per timepoint (wildtype), n=7-15 (Lmx1a KO). **H)** Predicted (line) and measured (dots) pSmad1/5 and Lmx1a degradation dynamics upon inhibition with 3μM LDN193189 from 72h after BMP addition (here designated as t=0h). n=8-12 (pSmad1/5), n=5-15 (Lmx1a) images per timepoint. Error bars, SD. **I)** Types of pSmad1/5 and Lmx1a dynamic behaviours (right) as a function of 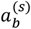 and 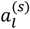. Star, parameters used in the model. Details in SI.

Numerical simulations indicated that this minimal circuit (Fig. 4A) produces a gradient of pSmad1/5 activity from the margin to the centre of colonies (SI) with an amplitude that overshoots before reaching steady-state pSmad1/5 levels at long times (Fig. 4B). Given that the pSmad1/5 gradient decays with near-constant length scale over time, consistent with edge activation and diffusion (SI), the temporal dynamics of the circuit can be captured by plotting the maximum levels of pSmad1/5 signalling at every time point (Fig. 4B). We therefore use this representation in subsequent analysis. Altogether, these simulations show that this simple negative feedback circuit qualitatively recapitulates the first phase of the temporal dynamics of pSmad1/5 signalling (Fig. 4B, 2C).

We then asked if this circuit alone could lead to oscillations in signalling that could potentially explain how pSmad1/5 is upregulated again in phase two. Analysis of the dynamics of the model as function of parameters and initial conditions showed that pSmad1/5 levels relax to either zero or non-zero steadystate level (Fig. 4C, SI). The downregulation of pSmad1/5 from its maximum to its steady state value differed across this parameter space (Fig. 4C). This allowed us to constrain the model parameters to the non-zero steady state regime and determine that the initial exogenous concentration of 0.5ng/ml BMP4 is above the steady-state levels of BMP (Fig. 4C, SI). Notably, this simple circuit does not predict a second large-amplitude pulse of pSmad1/5 activity in any parameter regime. This indicates that the minimum model (Fig. 4A) cannot explain the upregulation of pSmad1/5 in phase two.

The observation that BMP signalling in the initial phase is required for Smad1/5 signalling in the second phase (Fig. 3C, D) suggested the presence of a self-activating feedback loop on BMP acting at slower time scales compared to phase one. We hypothesised that such a positive feedback could be mediated via the formation of roof plate cells, which are a known source of BMP ligand expression in vivo ^11^. Lmx1a is an essential regulator of roof plate formation in vivo ^32^ and is itself a BMP target gene ^11^, therefore we included it as a candidate mediator of the positive feedback (Fig. 4E, SI).

If Lmx1a is required to upregulate pSmad1/5 in phase two, our model predicts that the absence of Lmx1a should significantly reduce, but not completely abolish, pSmad1/5 signalling at long times, while leaving the short time scale dynamics unchanged (Fig. 4F, SM). To test this, we generated Lmx1a knockout (KO) cells using CRISPR/Cas9 (Methods). Consistent with model predictions, in the Lmx1a KO cells, the pSmad1/5 levels were similar to control until 24h, but were strongly downregulated at later time points (Fig. 4D, G). To further validate the role of Lmx1a in upregulating pSmad1/5, we assessed the endogenous expression of BMP ligands in Lmx1a KO cells in phase two. This revealed that the expression of BMP6, BMP7 and GDF7 is strongly reduced in Lmx1a KO cells compared to control (Fig. S3D). These observations indicate that the second phase of BMP signalling depends on the induction of BMP ligand expression by Lmx1a, consistent with Lmx1a mutant phenotype in vivo ^33^.

To produce well-defined first and second phases in the model, we found that the characteristic time scale of pSmad1/5 activity had to be shorter than that of Lmx1a activity. This implies that the degradation rate of Lmx1a must be much lower than that of pSmad1/5. To test this, we measured the levels of pSmad1/5 and Lmx1a expression in colonies where BMP signalling was abruptly inhibited at t=72h using a high concentration (3μM) of the BMP inhibitor LDN193189 (Fig. 4H). Consistent with our prediction, these experiments indicated that the half-life of Lmx1a corresponds to ∼8h, while that of pSmad1/5 corresponds to ∼0.5h (Fig. 4H). We used these values to constrain the time scales of pSmad1/5 and Lmx1a dynamics. Based on this, simulations mimicking BMP signalling inhibition could quantitatively capture the experimentally observed decay dynamics (Fig. 4H).

Based on these inferred time scales, we found that the model captures the first and second phase separation in a configuration such that Lmx1a is weakly activated by pSmad1/5 (Fig. 4I), but beyond a threshold exhibits a pSmad1/5-dependent auto-activation (SI). This extended model captures a rapid initial phase driven by amplification of the exogenous BMP initially present in the system, followed by inhibitor-driven downregulation and subsequent increase in pSmad1/5 levels mediated by Lmx1a (Fig. 4F, G). To verify that Lmx1a self-activation occurs, we overexpressed Lmx1a using a doxycycline-inducible promoter (Methods). Consistent with the role of Lmx1a in inducing BMP production, we found that overexpression of Lmx1a increased the pSmad1/5 levels of BMP4 treated cells (Methods, Fig. S3E). Crucially, we found that in the absence of exogenous BMP4, Lmx1a overexpressing cells promoted endogenous Lmx1a expression in a non-cell autonomous manner, which is consistent with the Lmx1a self-activation suggested by the model (Fig. S3F).

Altogether, our analysis suggests that the two phases of pSmad1/5 signalling arise due to a time scale separation between two interacting subnetworks: a rapid first phase driven by a negative feedback loop on BMP levels by the induction of BMP inhibitors; and a slow second phase driven by a positive feedback loop that involves the transcription factor Lmx1a. The signal is relayed over time from the fast to the slow subnetwork.

### The initial signalling response is BMP concentration-sensitive but with a robust duration

Our model makes several experimentally testable predictions. First, it predicts that BMP inhibitors that temporally restrict the first phase of signalling are produced in a BMP-dependent manner (SI). To test the contribution of self-generated BMP inhibition to pSmad1/5 dynamics, we investigated which BMP signalling inhibitors might be responsible for the downregulation of pSmad1/5 in our system. RNA sequencing analysis indicated that BMP4 treatment induces the expression of a subset of several BMP inhibitors, including Noggin, Smad6 and Smad7 rapidly, at 8h (Fig. S4A). Treatment of the colonies with different BMP4 concentrations indicated that inhibitor induction occurs in a concentration-dependent manner, consistent with our model (Fig. S4A). HCR in situ hybridization further confirmed the upregulation of *Nog, Smad6* and *Smad7* in response to BMP4 (Fig. S4B). The expression of these BMP inhibitors was further maintained at 24h as pSmad1/5 levels are downregulated, which suggests that they are plausible mediators of the pSmad1/5 dynamics in this system.

To test the effects of these candidate inhibitors further, we used CRISPR/Cas9 mediated genome editing to generate knockout (KO) ES cell lines (Methods). Analysis of Nog KO cells revealed that the peak of pSmad1/5 signalling at 8h was similar to that in wildtype cells, while the subsequent downregulation occurred less efficiently in the knockout resulting in higher pSmad1/5 levels from 24h onwards compared to control cells (Fig. 5A, B). Similar results were obtained in cells with a double knockout of Smad6 and Smad7 (Smad6/7 DKO) (Fig. 5B, S5A). This phenotype is captured by our model with reduced production of BMP inhibitor (Fig. 5C). Together, these results indicate that the duration of the initial pSmad1/5 signalling phase depends on the pSmad1/5-dependent expression of several BMP inhibitors.

**Figure 5.**
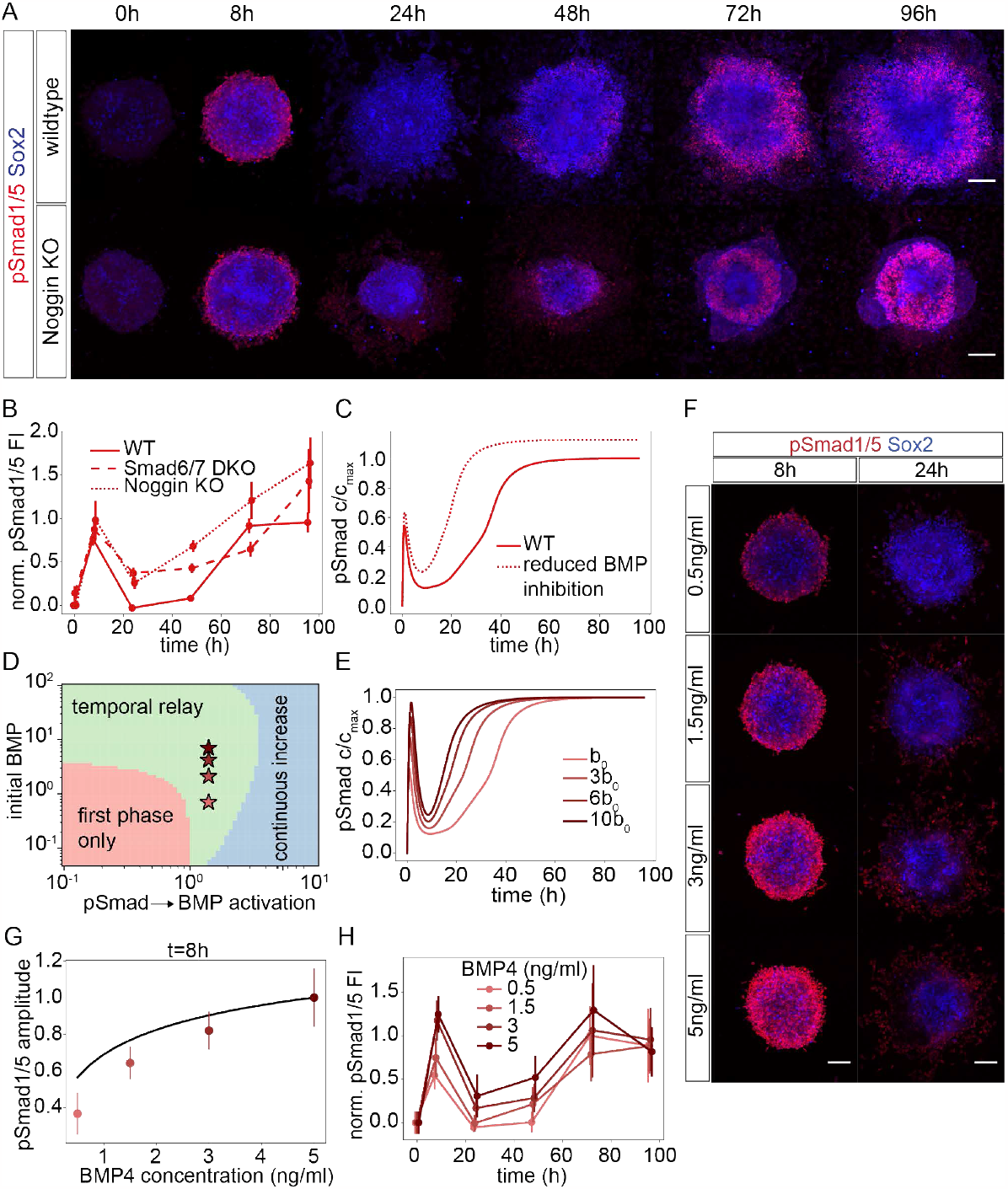
Signalling dynamics in BMP inhibitor knockouts and different concentrations of BMP4. **A**) Immunostaining against pSmad1/5 in wildtype and NogKO cells treated with 0.5ng/ml BMP4. Scale bar 100µm. **B)** Mean and 95% CI of pSmad1/5 fluorescence intensity (FI) in immunostainings. Images per timepoint from >2 experiments: n=43-97 (wt), n=23-39 (Noggin KO), n=12-18 (Smad6/7 DKO). **C)** Simulation of pSmad1/5 dynamics in control (solid) vs NogKO cells (dashed) by reducing the activation rate of the BMP inhibitor by 30%. **D)** Phase diagram of model behaviours (illustrated in Fig. 4I) as a function of 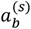 and *b*_0_. Stars, model parameters corresponding to BMP4 concentrations tested experimentally. **E)** Simulations of pSmad1/5 dynamics for 1,3, 6, and 10 times the initial concentration of BMP (*b*_0_) as indicated. **F)** pSmad1/5 and Sox2 immunostaining of cells treated with indicated BMP4 concentrations. Scale bar 100µm. **G)** Predicted (black line) and measured (dots) pSmad1/5 level at 8h. Error bars, SD. n=24-48 colonies per condition from 2 experiments. **H)** Quantification of pSmad1/5 FI from immunostainings for the indicated concentrations of BMP4. FI normalized to the max FI in the 0.5ng/ml BMP4 condition (Methods). n=15 images per timepoint and concentration. Mean with 95% CI is shown.

Another prediction of the model is that the biphasic dynamics are observed for a defined broad range of initial BMP concentrations (Fig. 5D, SI). Furthermore, the amplitude of the pSmad1/5 peak in phase one increases with increasing initial BMP concentration, which represents the exogenous BMP added to the culture media (Fig. 5E). Consistent with this prediction, the amplitude of the pSmad1/5 gradients in the differentiated colonies 8h after BMP treatment increases with the exogenous concentration of BMP4 in close quantitative agreement with the model (Fig. 5F-H, Fig. S5B). In addition, upon downregulation, pSmad1/5 reaches minimum levels that correlate with the initial concentration, yet the time of the downregulation is robust to the concentration of BMP in both the model and the experimental data (Fig. 5F, S5C, SI). This shows that the duration of the first phase of pSmad1/5 signalling is independent of the exogenous BMP concentration. Our model further indicates that in contrast to the first phase, the maximum levels reached during the second phase are independent of the initial BMP concentration (Fig. 5E). Thus, at long times, the system approaches a steady state which is independent of the initial conditions. The slightly faster approach to steady-state in the model compared to experimental data is likely due to additional interactions of Lmx1a that are not incorporated in the model. Nevertheless, the model indicates that the time scale of the approach to steady state is faster for higher concentrations, which is consistent with the differences observed in the discrete time points (between 48h and 72h) sampled in our experimental data (Fig. 5H).

Together, our analysis shows that the initial peak pSmad1/5 amplitude provides a rapid and sensitive readout of the exogenous BMP concentration, whereas the underlying negative feedback mechanism via the BMP inhibitors ensures the robust termination of the first phase of pSmad1/5 signalling. Finally, the second phase converges to a concentration-independent pSmad1/5 signalling level which depends on Lmx1a.

### The biphasic BMP signalling dynamics influence pattern formation via Lmx1a

How do the dynamics of BMP signalling influence downstream pattern formation? To address this, we first asked what is the effect of reduced BMP inhibitor production on the expression of Lmx1a itself. The model predicts that this will lead to an earlier increase and higher levels of Lmx1a expression (Fig. 6A). Consistent with this, Lmx1a is upregulated in Nog KO cells compared to wildtype controls, and is also observed at earlier time points (Fig. 6B, C). Similarly, Smad6/7 DKO cells have increased Lmx1a levels (Fig. 6B, C). These results show that the efficient downregulation of pSmad1/5 after the first phase prevents premature Lmx1a increase and thereby limits Lmx1a expression.

**Figure 6.**
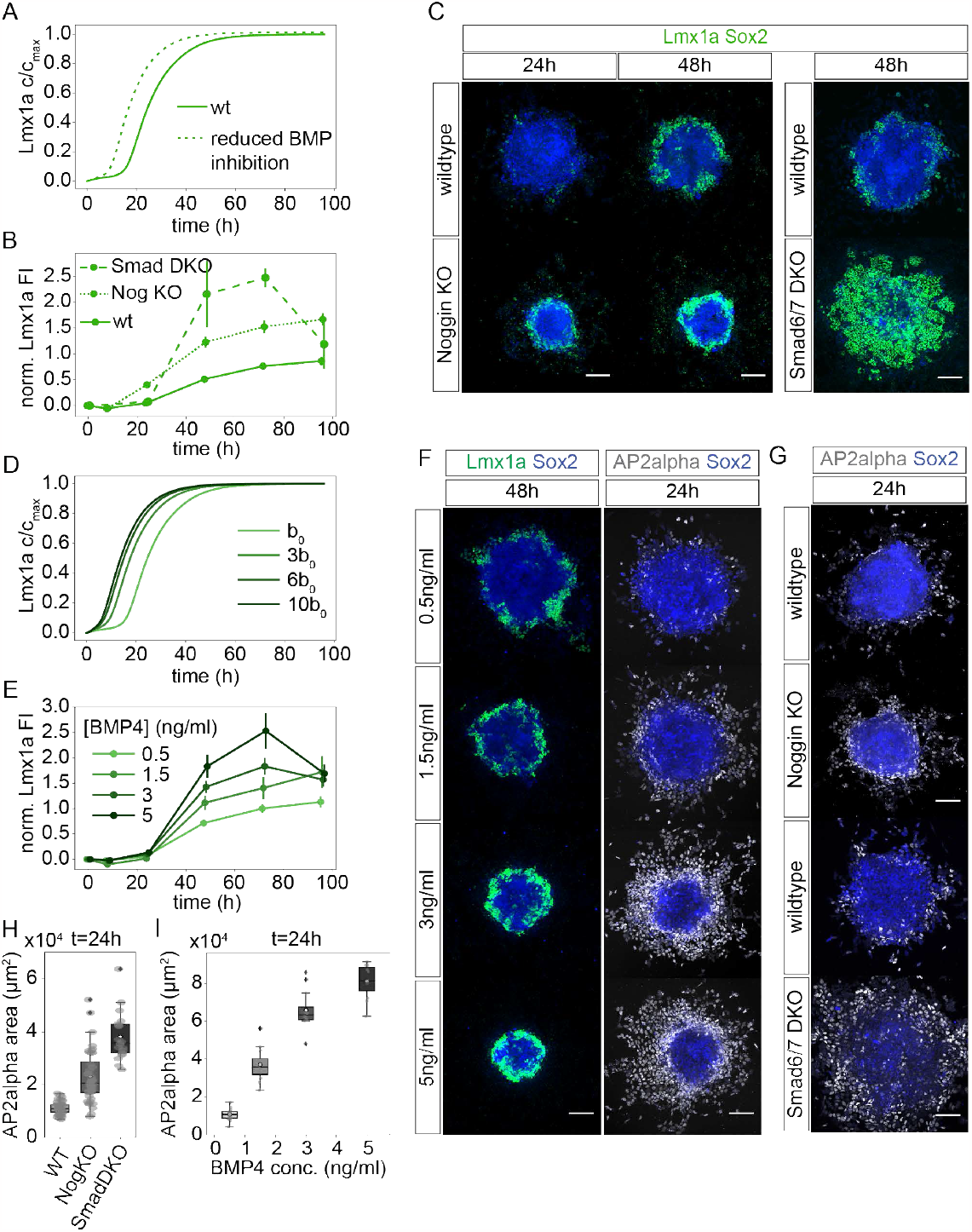
Lmx1a and neural crest dynamics upon perturbations in BMP signalling. **A**) Simulation of Lmx1a levels for wt (solid) and 30% reduced BMP inhibitor activation rate (dotted). **B)** Quantification of maximum Lmx1a FI from immunostainings. Images per time point: n=7-55 (wt), n=8-34 (NogKO), n=3-7 (Smad6/7 DKO). Mean and 95% CI are shown. **C)** Lmx1a immunostainings in wildtype, Nog KO and Smad6/7 DKO cells. Scale bars 100µm. **D)** Simulations of Lmx1a dynamics for 1,3, 6, and 10 times the initial concentration of BMP (b0) as indicated. **E)** Quantification of Lmx1a (as in B) for wildtype cells cultured at the indicated concentrations of BMP4. FI normalized to 72h 0.5ng/ml condition (Methods). n=5-30 colonies per data point. **F)** Colonies treated with indicated BMP4 concentrations stained for Lmx1a and AP2alpha. **G)** Immunostaining against AP2alpha in wildtype, Nog KO and Smad6/7 DKO cells treated with 0.5ng/ml BMP4. **H)** Quantification of AP2alpha expression area in the indicated genotypes at 24h. n=28 (wt), n=24 (NogKO), n=18 (SmadDKO). **I)** AP2alpha expression area at 24h correlates with BMP4 concentration. n=12 colonies per concentration. I, H: Box, median and IQR; white circle, mean.

Our experimental and theoretical results imply that Lmx1a expression is dependent on the first BMP signalling phase. Consistent with the sensitivity of the initial pSmad1/5 amplitude to the exogenous BMP4 concentration (Fig. 5F-H), the pre-steady state levels of Lmx1a are also predicted to be dependent on the initial BMP4 concentration (Fig. 6D). The experiments confirm this prediction: the level of Lmx1a expression increases with BMP4 concentration prior to 96h (Fig. 6E, F). Consistent with the concentration-independent steady state predicted by the model, the Lmx1a levels observed at last time point in our time course (96h) converge to a common state (Fig. 6E). These observations indicate that the levels of BMP signalling begin to influence the amount of Lmx1a activation during the initial phase.

The initial phase of BMP signalling is also critical for the specification of neural crest. AP2alpha and Sox9 expression are first detected at low levels at 8h and migrating neural crest cells outside of the Sox2 colony are visible at 24h (Fig. 1D, F). Similar to Lmx1a, the amount of AP2alpha expressing neural crest cells is significantly increased after 24h in Nog KO and Smad6/7 DKO cells, which have extended duration of signalling, compared to control cells (Fig. 6G, H). Furthermore, the amount of AP2alpha cells at 24h increases in proportion to the concentration of exogenous BMP4 and the amplitude of the pSmad1/5 gradient at 8h (Fig. 6F, I). Neural crest cells lose Sox2 expression as they undergo epithelial to mesenchymal transition and migrate out of the colony, which results in smaller Sox2+ colonies at higher exogenous BMP4 concentrations (Fig. 6F). Overall, similar to Lmx1a, these observations indicate that both the levels of pSmad1/5 signalling, starting from the initial phase, as well as the duration of signalling, affect the amount of neural crest that it specified.

To rule out that the changes in Lmx1a dynamics that we observe are an indirect consequence of neural crest specification, we inspected Lmx1a domain formation in Sox9 knockout cells (Methods). Sox9 mutant mouse embryos lack functional neural crest due to a defect in neural crest survival ^34^. Consistent with the mouse phenotype, the number of AP2alpha neural crest cells is markedly reduced in Sox9 KO cells (Fig. S5D). Nevertheless, the onset and levels of Lmx1a expression in Sox9 KO is similar to control cells (Fig. 7A, B). This is consistent with previous qualitative observations of Sox9-/-embryos ^34^ and suggests that NC does not affect the specification of roof plate fate.

**Figure 7.**
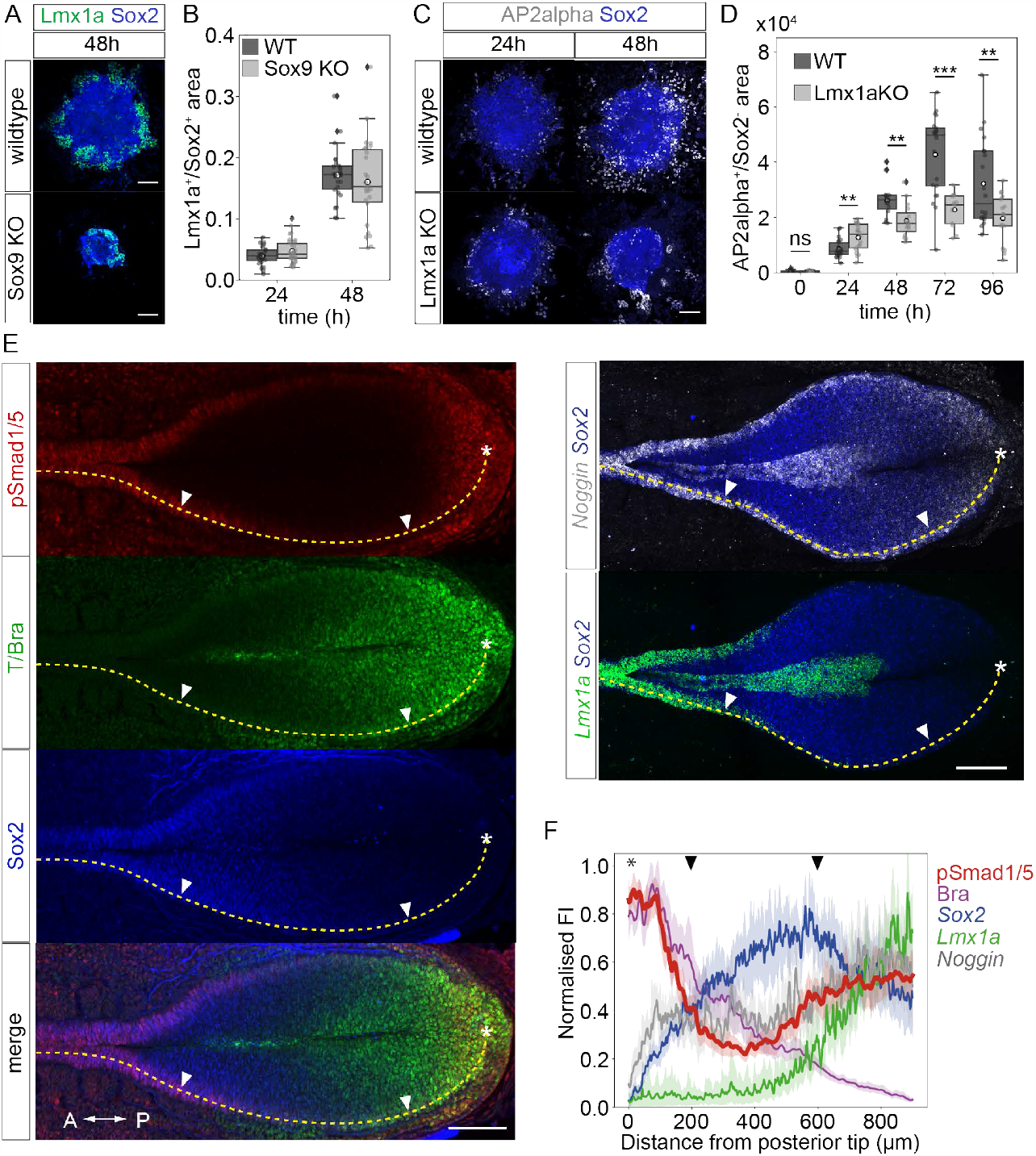
pSmad1/5 signalling dynamics in the posterior mouse spinal cord at E8.5. **A**) Wildtype and Sox9 KO cells stained for Lmx1a. **B)** The Lmx1a+ relative to total Sox2 expression area is similar in wildtype and Sox9 KO cells. n=31-36 (wt), n=31-44 (Sox9 KO) colonies per timepoint. In B, D: box median and IQR, white circle mean. **C)** AP2alpha levels are increased at 24h but decreased at 48h in Lmx1a KO cells compared to control. **D)** Quantification of AP2alpha^+^ Sox2 negative expression area in wildtype and Lmx1a KO cells. n=5-17 (wt), n=5-15 (Lmx1a KO). Two-tailed t-test p-values: **p≤0.005, ***p≤0.0005. **E)** Maximum FI projections of immunostainings against pSmad1/5, T/Bra, Sox2 and HCR against *Noggin, Lmx1a, Sox2* in the neural plate of E8.5 mouse embryos. Anterior, left. **F)** Immunostaining and HCR FI profiles measured along the border of the Sox2 expression domain (dashed line) from the posterior tip of the embryo (*). Arrows indicate region where BMP signalling levels are downregulated. n=10-14 profiles from 5-7 embryos.

To assess the effects of the second signalling phase and the roof plate, we inspected pattern formation in the Lmx1a KO cells. The formation of neural progenitor domain dp1, characterised by Atoh1 expression, was impaired in the Lmx1a KO (Fig. S5E), consistent with the in vivo phenotype of mutants that lack roof plate ^35^. By contrast, during the early stages of differentiation, at 24h, Lmx1a KO colonies had increased levels of AP2alpha cells (Fig. 7C, D). This was surprising and suggested that at this early stage Lmx1a has an inhibitory effect on neural crest specification. Later on, at 72h and 96h, neither AP2alpha nor Sox9+ neural crest cells could be observed outside the Sox2+ core of the colony in Lmx1a KO cells, indicating that Lmx1a is required for the normal maturation of neural crest, either directly or indirectly via signals produced by Lmx1a cells. Thus, the phenotype of Lmx1a KO cells in our differentiation protocol suggests that Lmx1a performs a dual role – on one hand, it inhibits the early neural crest specification, and on the other hand, it promotes the maturation and migration of committed neural crest cells. Altogether, our findings indicate that both the amplitude and duration of the early signalling phase are critical for the correct balance between neural crest, Lmx1a domain and neural domain (marked by Sox2). In turn, the second signalling phase is required for the maturation and maintenance of neural crest, roof plate expansion and the specification of neural progenitor subtypes.

### Biphasic BMP signalling dynamics underlie pattern formation in vivo

The observation that BMP signalling dynamics are biphasic in vitro, raises the question whether this is also observed in the developing mouse embryo. To address this, we immunostained E8.5 embryos for pSmad1/5, as well as Sox2 and Bra, which are co-expressed in neuromesodermal progenitor cells. Over time, the embryo undergoes posterior extension and cells shift their mean relative positions away from the posterior tip of the embryo. Although measurements of the exact cell trajectories during this process are not available, the spatial profiles of gene expression and signalling along the anterioposterior (AP) axis represent approximately the temporal changes experienced by cells. We therefore quantified the levels of pSmad1/5 signalling along the AP axis, focusing on the neural plate border region which most closely corresponds to the outer edge of in vitro generated colonies. Notably, we observed that along the border of the caudal neural plate, the levels of pSmad1/5 signalling are not uniform. Instead pSmad1/5 levels are high within and posterior to the caudolateral epiblast region where NMPs reside, subsequently decline along the posterior caudal neural plate border, and increase again in the anterior caudal neural plate (Fig. 7E, F). Furthermore, similar to the in vitro culture, we detected *Noggin* transcript expression (Methods) in the region where pSmad1/5 levels were low (Fig. 7E-F). In addition, Lmx1a expression was detected in the anterior caudal neural plate, at AP positions that coincided with increased pSmad1/5 levels (Fig. 7E-F). These observations suggest that similar to the in vitro situation, the levels of pSmad1/5 experienced by cells at the neural plate border undergo biphasic dynamics, in which the second phase of signalling occurs as Lmx1a expression becomes markedly upregulated.

To test whether BMP signalling dynamics affect pattern formation in vivo as they do in vitro, we analysed the effects of genetic deletions of BMP inhibitors on Lmx1a at different time points. Our model predicts that early deletion of BMP inhibitors will lead to a pronounced increase in levels of pSmad1/5 and Lmx1a expression, while later deletions will have a lesser effect (Fig. S6C, SI). To test this, we conditionally deleted Noggin in mouse embryos using a floxed allele (Methods) crossed to two different Cre lines (Fig. S6D, E). In the first case, Nog was knocked out in Sox2+ cells using tamoxifen-inducible Sox2-CreERT2 induced at E6.5. Analysis of Lmx1a expression in these animals at E10.5 of development indicated that they have larger roof plate compared to controls (Fig. S6D), consistent with the in vitro observations in Nog KO cells (Fig. 6B, C). In the second case, Nog was specifically deleted in Wnt1 expressing cells using Wnt1-Cre (Methods). The endogenous expression of Wnt1 is initiated in the closed neural tube (Fig. S6F), hence this represents a later deletion of Nog. In contrast to the early deletion, the RP size in Wnt1-Cre::Nog^Flox/Flox^ mutants was similar to control littermates (Fig. S6E).

Altogether, these results indicate that pSmad1/5 levels are temporally regulated in the neural plate by BMP inhibitor expression prior to the endogenous expression of Wnt1, which marks the second phase of BMP signalling in vitro. The downregulation of pSmad1/5 in the neural plate border restricts the duration of pSmad1/5 signalling and thereby influences the size of the Lmx1a domain. Altogether, our findings reveal how roof plate formation and dorsoventral patterning of the closed neural tube become dependent on earlier BMP signalling that occurs as cells leave the caudolateral epiblast.

## Discussion

Signalling relays have long been considered as possible mechanisms of morphogen gradient formation^36^. In these mechanisms, morphogens that spread at short range expand their activity range by inducing ligand production in adjacent cells. Recently, examples of this have been uncovered in the context of Nodal signalling during zebrafish and human gastrulation ^37,38^. Relay mechanisms that can give rise to millimetre long gradient ranges, such as the Wnt signalling gradient in planaria, have also been described ^39^. Here we uncovered a new type of a relay mechanism that intrinsically operates over time, rather than in space, creating sequential induction of morphogen ligand production and thereby signalling phases. This enables morphogen reuse within a tissue for different purposes – first to rapidly specify the neural plate border in a concentration-sensitive manner, and then to reform the gradient and use it to specify the fates of dorsal neural progenitor subtypes along the DV axis.

The mechanisms that contribute to dynamic changes in signalling can be difficult to manipulate and disentangle in vivo. We circumvented this challenge by developing a 2D in vitro model of the dorsal neural tube. 2D in vitro systems have previously been used to study signalling and pattern formation in mouse and human gastrulation ^20–22,28,40^. In contrast to these systems, our system represents a later developmental stage, early neurulation, and focuses on the posterior NMP-derived fates, rather than anterior patterning. Notably, unlike differentiation on restricted micropatterns, it does not limit growth and cell migration, and this was key for the formation of reproducible self-organised patterning. Given that pattern formation usually occurs in tight coordination with tissue growth, the stencil-based system that we developed could be widely applicable to other studies of pattern formation, signalling and growth control ^41^.

The stencil-based system allowed us to obtain quantitative readout of BMP signalling dynamics that we combined with biophysical modelling to identify the underlying mechanisms. Temporal adaptation in the continued presence of ligand, similar to the first phase of BMP signalling dynamics that we observe, can arise from diverse cellular mechanisms ^4,6,42,43^. Here we found that in dorsal neural progenitors, BMP signalling is downregulated by BMP-dependent induction of inhibitors. In contrast to other temporal adaptation mechanisms where the pathway remains refractory over long times, in the dorsal neural tube, the induction of a secondary network operating at slower time scales bypasses such adaptation and initiates a new phase of signalling. These connected networks ensure robust sequential relay of signalling in a wide region of parameter space (Fig. 4I, 5D and SI).

Our analysis indicates that the two subnetworks are coupled by the transcription factor Lmx1a. The second phase of signalling is initiated by the dependence of Lmx1a induction on a pSmad1/5 activation threshold in the first phase. This is supported by our observations that Lmx1a is expressed early on, at 24h in vitro, and in the open neural plate, and by single cell RNA-sequencing data indicating its expression in cells with early neural plate border signature ^44^. Previous studies of Lmx1a have used the Dreher (dr^J^) mouse mutant ^45^, in which Lmx1a is initially normally expressed in the neural plate at E8.5, but progressively lost in the closed neural tube ^33^. The Lmx1aKO cells that we generated thus helped to identify and validate a previously unrecognised early function of Lmx1a in coordinating the biphasic pSmad1/5 dynamics.

An intrinsic property of the temporal relay mechanism that we uncovered is that is serves as a checkpoint between signalling events, thereby enforcing a temporal order of downstream patterning events. As cells exit the neuromesodermal progenitor state, they respond to BMP signalling to initiate neural crest specification, consistent with previous studies ^46^. Continued neural crest development requires a second phase of signalling ^16,47^. At the same time, neural crest specification is followed by roof plate-dependent patterning of the dorsal neural progenitor subtypes ^35^. The temporal relay ensures this order and provides a potential mechanism for timing the transition between the neural plate border phase and the subsequent roof-plate dependent patterning phase.

The relay mechanism allows to robustly control distinct features of the signalling dynamics during different phases. The initial phase and concomitant specification of neural crest are BMP concentration-sensitive. This is in agreement with previous studies that found that the formation of non-neural ectoderm, neural plate border and neural ectoderm occurs in response to a concentration gradient of BMP ^48–50^. In this context, BMP inhibition is thought to be required to achieve the lowest levels of BMP signalling necessary for the acquisition of neural ectoderm identity. By contrast, we show that BMP inhibitors primarily limit the duration of the initial concentration-sensitive phase and thereby the amount of neural crest. Our analysis further indicates that during the second signalling phase, the system responds to different BMP concentrations by slightly altering the timing of the response. This could provide a potential explanation for the observations that both the concentration and duration of BMP signalling affect the acquisition of specific dorsal neural progenitor identities ^9,12^.

Similar to other studies, we also show that BMP signalling is required to initiate the specification of neural plate border during a distinct time window at the end of gastrulation ^51–53^. However, our study leaves open the question of what regulates the requirement for cells to respond to BMP signalling immediately after exiting the NMP state and not later (Fig. 3A). Previous studies have suggested that the competence to specify neural crest depends on the repression by Fgf signalling of neural progenitor specific transcription factors, such as Pax6, that inhibit NC specification ^53^. Such a mechanism could operate in parallel with the temporal relay in BMP signalling that we describe.

In summary, while the temporal dynamics of morphogen signalling influence cell fates specification in multiple systems ^1–3^, the underlying mechanisms are in many cases still unclear. This study uncovers a mechanism by which morphogen signalling is relayed over time in the dorsal neural tube. A temporal relay mechanism might also operate in the ventral neural tube. Similar to the dorsal neural tube, there, pattern formation is initiated by Shh produced in the notochord, and subsequently depends on the floor plate. A related mechanism in which BMP4 ligand production is regulated by positive feedback has also been observed during exit of ES cells from pluripotency and commitment to differentiation ^54^. Thus, temporal relay mechanisms may represent a general strategy to time developmental transitions.

## Supplementary Figures S1-S6

**Figure S1.**
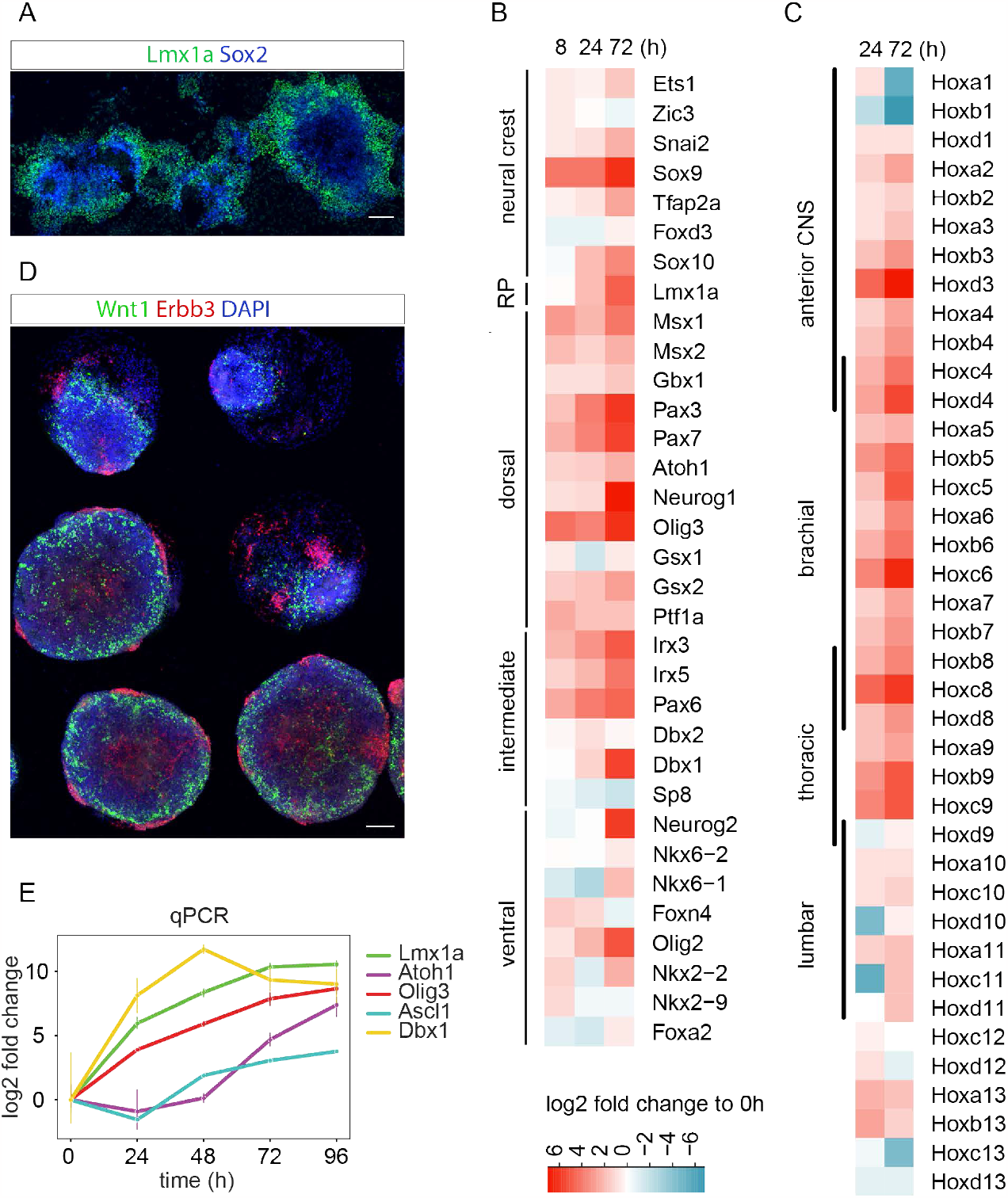
Dorsoventral and anterioposterior identities of differentiated cells. **A**) Radial gene expression pattern in colonies cultured without stencils. Scale bar 100µm. B) Expression of typical DV identity genes over time relative to 0h from RNA-sequencing experiments. In B, C: n=3 samples per time point. C) Expression of Hox genes over time relative to 0h from RNA-sequencing. D) Example colonies cultured on confined micropatterned surfaces. Immunostaining against Wnt1 (RP) and Erbb3 (NC). Scale bar 100µm. E) Expression of neural progenitor subtype markers relative to 0h measured by qPCR. n=1-10 samples per time point per gene. Error bars, SD.

**Figure S2.**
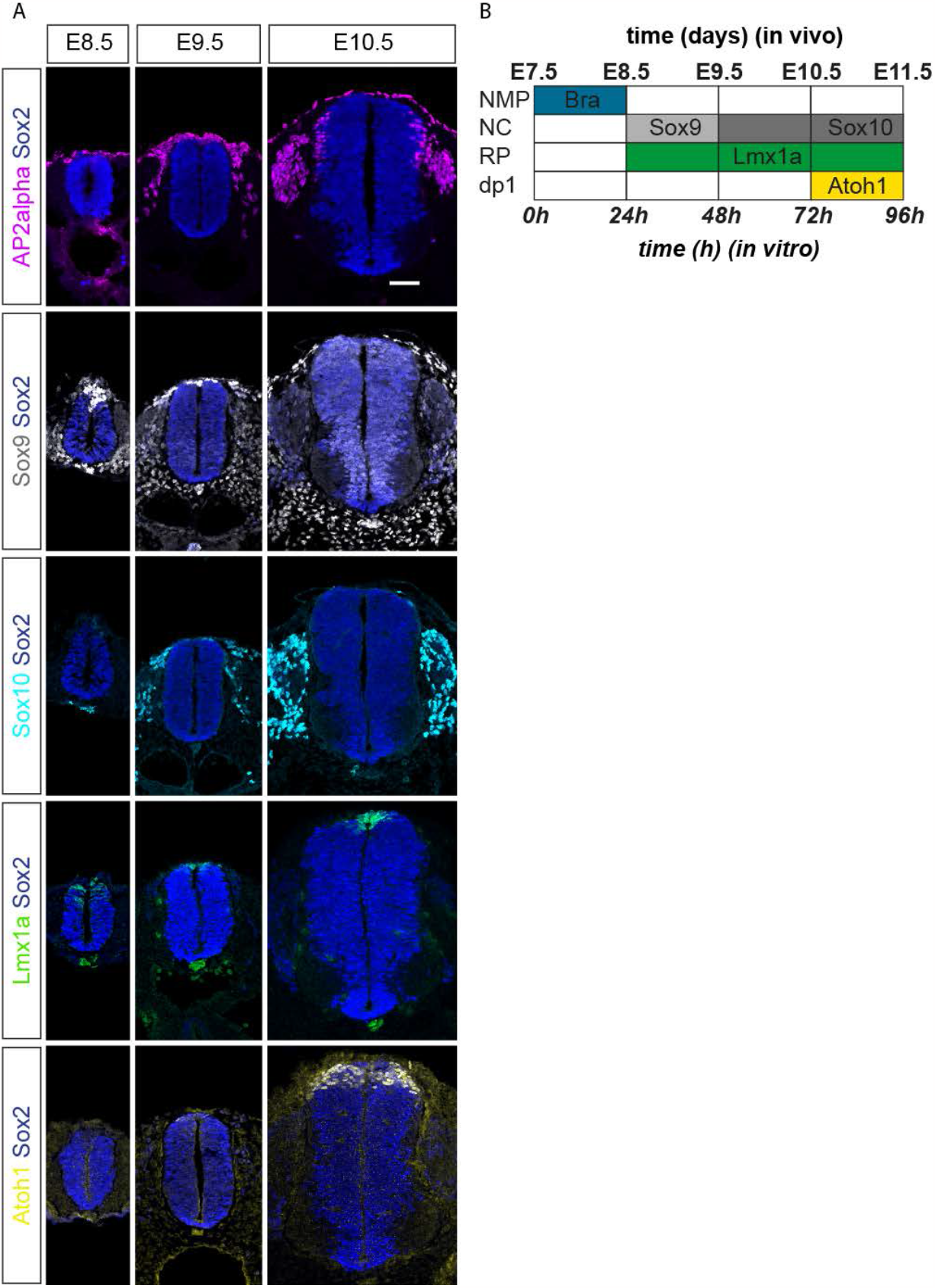
Spatiotemporal sequence of dorsal neural tube patterning in vivo. **A**) Immunostainings of transcription factors expressed in NC (AP2a, Sox9, Sox10), RP (Lmx1a) and dP1 (Atoh1) domains at the indicated stages of mouse development. Scale bar 50µm. B) Time of expression of genes characteristic for NMPs and NC, RP and dP1 progenitors in the mouse neural tube (embryonic days indicated above) and in vitro (hours post BMP addition indicated below).

**Figure S3.**
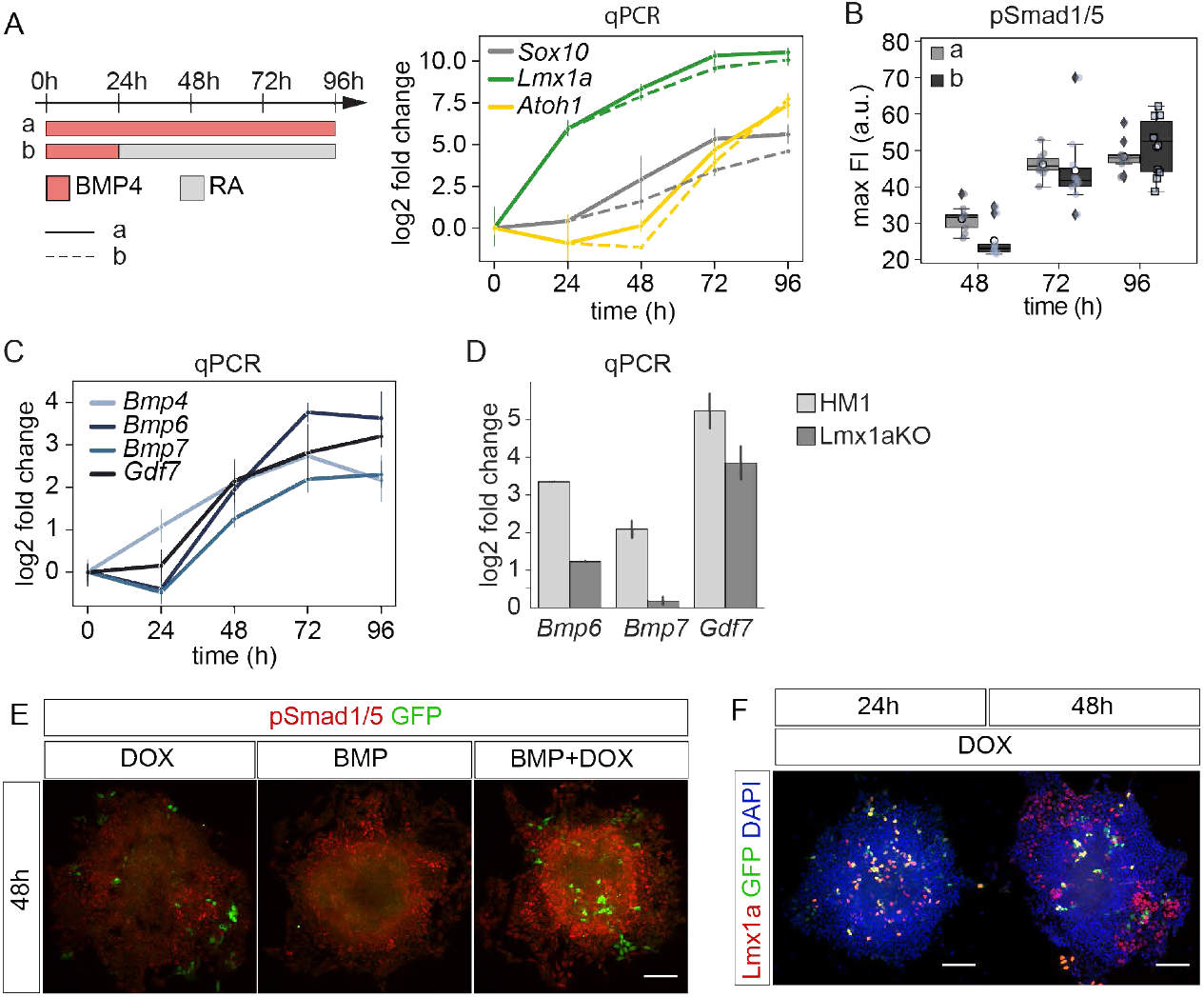
The second signalling phase results from Lmx1a self-activation and endogenous BMP ligand expression. **A**) Schematic of experimental conditions (left). Continuous treatment (a, solid) or 24h pulse of 0.5ng/ml BMP4 (b, dashed). Expression of indicated genes relative to 0h measured by qPCR. n=1-8 samples per time point per gene. Error bars, SD. **B)** Quantification of pSmad1/5 FI from immunostainings for the conditions in A. n=9-10 per time point per treatment. Box median and IQR, white circle mean. Twotailed T-test, p< 0.05 for t=48h, ns for 72h and 96h. **C)** Expression of indicated BMP ligands relative to 0h measured by qPCR in cells treated with 0.5ng/ml BMP4, n=2-6 samples per time point per gene, Error bars, SD. **D)** qPCR analysis of BMP ligand expression in wildtype and Lmx1a KO cells treated with 0.5ng/ml BMP4. T test, p< 0.001 for Bmp6, p< 0.05 for Bmp7, ns for Gdf7. n=2, Error bars, SD. **E)** pSmad1/5 and Lmx1a-IRES-GFP expression in TetON-Lmx1a-IRES-GFP cells treated with 0.5ng/ml BMP4, doxycycline, or 0.5ng/ml BMP4+ doxycycline as indicated. Scale bar 100µm. F) Lmx1a immunostaining (red) indicating endogenous Lmx1a expression and GFP expression (green) in TetON-Lmx1a-IRES-GFP cells treated with doxycycline. Scale bar 100µm.

**Figure S4.**
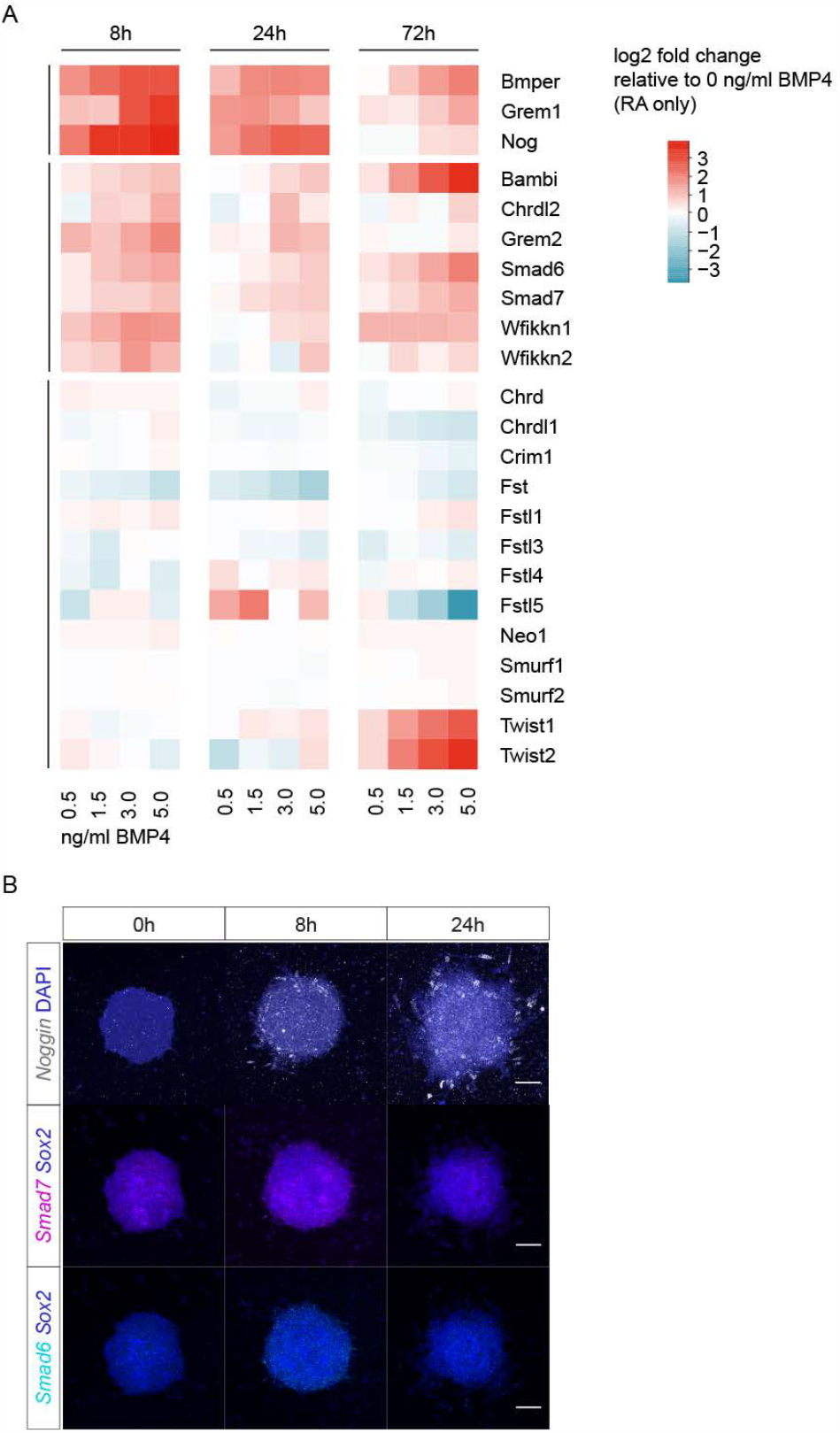
Expression of BMP pathway inhibitors upon treatment with different concentrations of BMP4. **A**) Expression levels of BMP inhibitors in cells treated with different BMP concentrations at the indicated time points relative to RA only (0 ng/ml BMP4) condition determined by RNA-sequencing. Rows were clustered by K-means clustering using the 8h time point (3 centres, black lines). n=3 samples per condition and timepoint. **B)** HCR in situ hybridisation shows spatial expression profiles of the indicated BMP inhibitors at the indicated time points. Scale bar 100µm.

**Figure S5.**
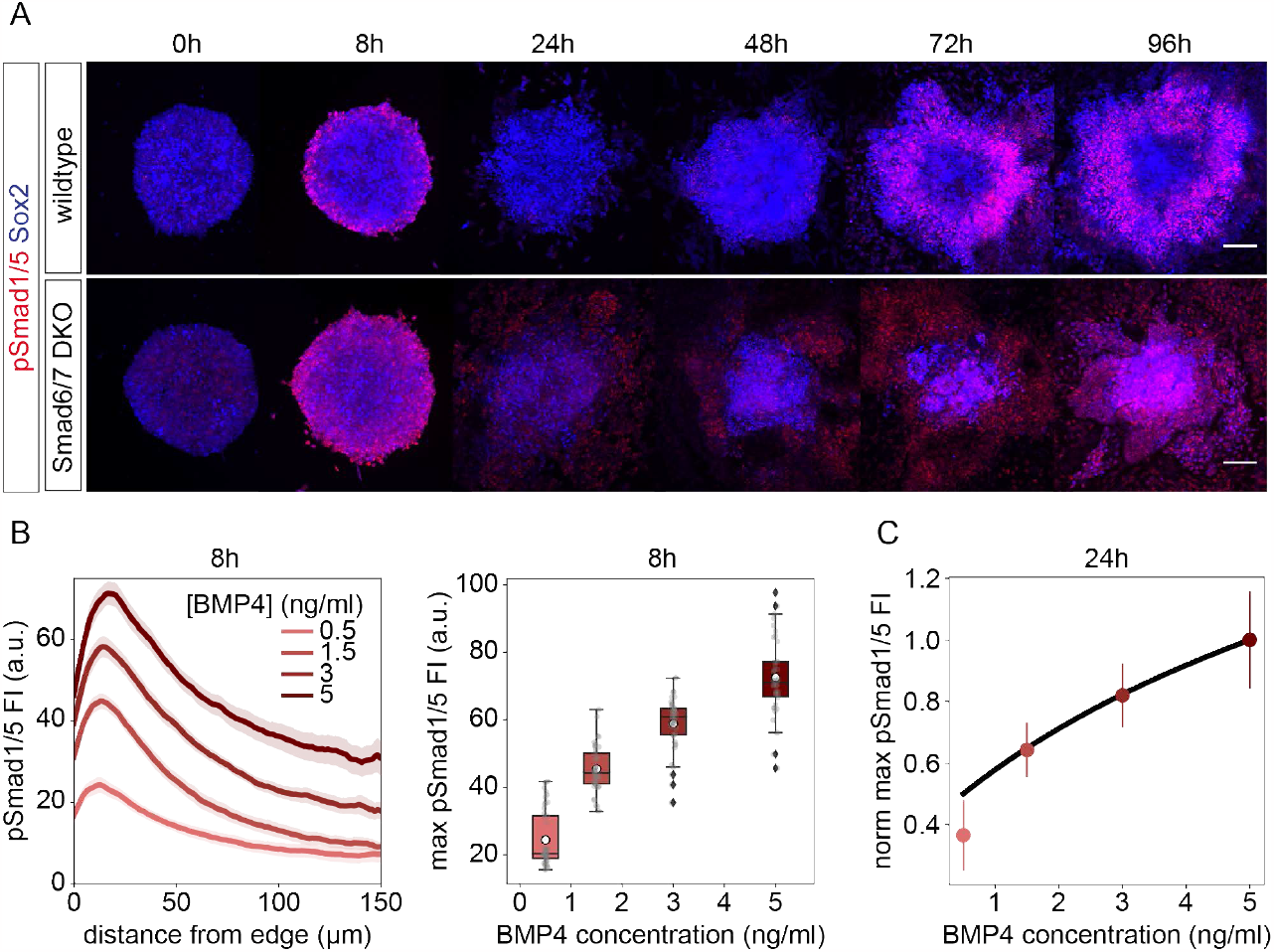
BMP signalling and target gene dynamics upon perturbations. **A**) Immunostaining against pSmad1/5 in wildtype and Smad6/7 DKO cells treated with 0.5ng/ml BMP4. Scale bar 100µm. **B)** Mean spatial profiles of pSmad1/5 FI at 8h for different concentrations of BMP4. Shaded regions, 95% CI (left). Maxima of the spatial profiles at 8h for different concentrations of BMP4 (right). n=24-48 samples per condition. **C)** Predicted (black line) and measured (dots) FI maxima of the mean pSmad1/5 spatial profiles at t=24h, corresponding to the time of pSmad downregulation after the first phase. FI values are normalised to the 5ng/ml BMP4 condition. n=17-26 samples per condition. Error bars, SD.

**Figure S6.**
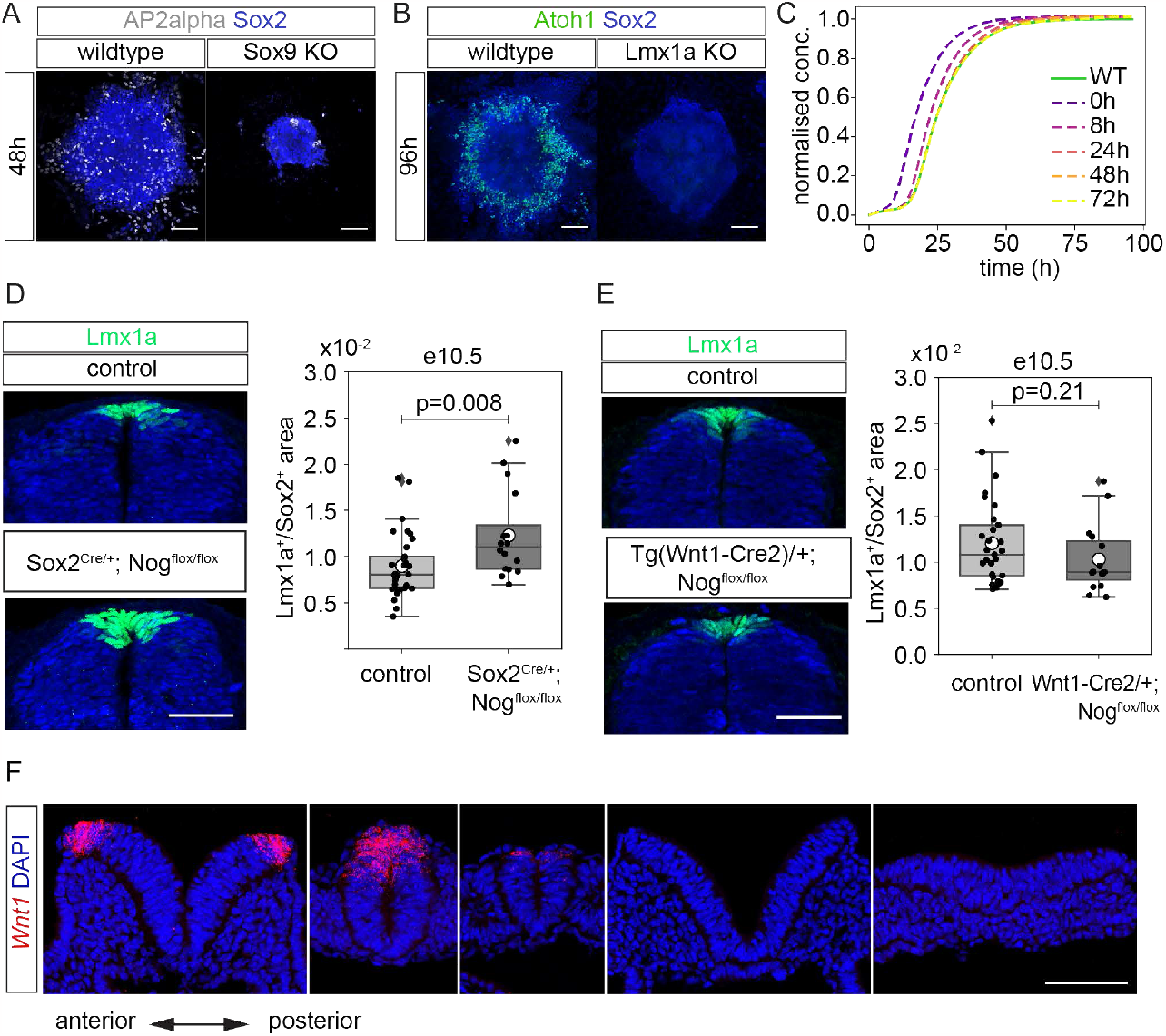
Lmx1a expression is most affected by early perturbations in BMP signalling. **(A)**AP2alpha expression is strongly decreased in Sox9 KO cells compared to wildtype. Scale bar 100µm.(B)Lmx1a KO cells do not express Atoh1 indicating absence of the dp1 domain. **C)** Simulations of Lmx1a levels for different time points of BMP inhibitor reduction. **D)** Sox2 (blue) and Lmx1a (green) immunostaining in brachial sections from control littermates and Sox2^CreER/+^; Nog^Flox/Flox^ mutants generated by tamoxifen injection at E6.5 and collected at E10.5. Scale bar 50µm (left). Quantification of the Lmx1a expression area relative to Sox2 (right). Sections from n=17 control and 8 KO embryos from 3 litters. **E)** Immunostaining and quantification as in D, but for Tg(Wnt1-Cre2)/+; Nog^Flox/Flox^ mutants and control littermates collected at E10.5. Sections from n=15 control and 6 KO embryos from 3 litters. Boxes in D, E show median and IQR. While circle indicates mean. Two-tailed T-test p-value indicated. **F)** HCR in situ hybridisation for Wnt1 in transverse mouse embryo sections at E8.5. Wnt1 is first detected in the closed neural tube, but not in the caudal neural plate. Scale bar 100µm.

## Materials and methods

### ES cell maintenance culture

Mouse HM1 ES cell ^55^ background was used for all experiments. All cell culture reagents and plastics are described in Tables T1 and T2, respectively. ES cells were maintained on CellBIND dishes (Corning) coated with 0.1% gelatin (Sigma) at 37°C 5% CO_2_.

**Table T1.** Media and Supplements.

| Name | Source | Cat. No. |
| --- | --- | --- |
| 0.1% gelatin in H<sub>2</sub>O | Millipore | ES-006-B |
| 2-Mercaptoethanol | Gibco | 31350010 |
| Accutase | Gibco | A1110501 |
| B-27 Supplement (50X) | Gibco | 17504001 |
| bFGF | R&D | 3139-FB-025 |
| BMP4 | R&D | 5020-BP-010 |
| BSA | Sigma-Aldrich | A3156 |
| CHIR99021 | Axon | 1386 |
| DMEM/F-12 | Gibco | 21331020 |
| Fetal bovine serum (FBS) for ESC (30 min. 56°C heat inactivated) | Pan Biotech | P30-2602 |
| Gelatine Solution, 2% in H<sub>2</sub>O | Sigma-Aldrich | G1393 |
| GlutaMAX | Gibco | 35050061 |
| L-Glutamine | Gibco | 25030024 |
| LDN-193189 | Tocris | 1517128 |
| LIF | Chemicon | ESG1107 |
| MEM Non-Essential Amino Acids Solution | Gibco | 11140035 |
| N-2 Supplement (100X) | Gibco | 17502001 |
| Neurobasal Medium | Gibco | 21103049 |
| PD98059 | Cell Signalling Technology | 9900 |
| Penicillin-Streptomycin | Gibco | 15140122 |
| Retinoic Acid (RA) | Sigma-Aldrich | R2625 |
| ROCK inhibitor | Tocris | 1254 |
| Doxycycline | Sigma | D9891 |

**Table T2.** Cell culture plastics.

| Name | Source | Cat. No. |
| --- | --- | --- |
| CellBIND surface, 60 mm | Corning | 3295 |
| CellBIND surface, 100 mm | Corning | 3296 |
| CellBIND surface, 12-well | Corning | 3336 |
| μ-slide 2-well, ibiTreat | ibidi | 80286 |
| CYTOOchips Arena A | Cytoo | 10-020-00-18 |

Stem cells were maintained in N2B27 + 2i + LIF media. N2B27 is composed of 1:1 mix of DMEM/F-12 (Gibco) and Neurobasal medium (Gibco), 1X N2 (Gibco), 1X B27 (Gibco), 0.08% BSA (Sigma), 2 mM L-Glutamine or Glutamax (Gibco), 100 U/ml Penicillin-Streptomycin (Gibco), 0.1 mM 2-Mercaptoethanol (Gibco). For 2i + LIF, N2B27 was supplemented with 3µM CHIR99021 (Axon), 1µM PD98059 (Cell Signalling Technology) and 1,000 U/ml LIF (Chemicon). Media was exchanged daily.

Cells were passaged every 2-3 days by incubation with Accutase (Gibco) for 2-3 min at 37°C, dissociated to a single cell suspension. Cells were counted using a cell counter (BioRad), collected by centrifugation at 1000rpm for 4 min, then seeded at a density of 120-200x10^3^ cells per 6 cm plate. Cell lines were routinely tested and confirmed negative for mycoplasma.

### Generation of knockout and Lmx1a overexpressing ES cell lines

To generate knockout cell lines, gRNAs were designed using the CRISPOR tool ^56^ against sequences in exons 1 to 3 (Table T5). Two gRNAs per gene with MIT score >70 were chosen, phosphorylated oligos ordered from Sigma and cloned into pSpCas9(BB)-2A-Puro (PX459) V2.0 (Addgene, Plasmid #62988) or pSpCas9(BB)-2A-GFP (PX458) (Addgene, Plasmid #48138) according to standard protocol ^57^.

To generate the TetON-Lmx1a-IRES-GFP cell line, the mouse Lmx1a CDS was amplified from cDNA. PB-TRE-EGFP-EF1a-rtTA (Addgene, Plasmid #104454) was digested with NheI and KpnI and the Lmx1a CDS and IRES-GFP fragments were inserted using the In-Fusion HD Cloning kit (Takara). Molecular biology kits used are listed in Table T4.

A total of 3-4µg of plasmid DNA was electroporated into 3-4x10^6^ early passage ES cells using the Amaxa Nucleofector II (Lonza) programme A-023. Electroporated cells were seeded onto a gelatine coated 10cm CellBind plate (Corning) and cultured in 2i+LIF. The following day, 2i+LIF was supplemented with 2µg/mL puromycin (Sigma) for 48 hours (or longer for the TetON-Lmx1a-IRES-GFP line), or sorted for GFP expression. After additional 5-7 days in 2i+LIF, individual colonies were picked into 96-well plates, dissociated using Accutase and plated onto feeder cells (mouse embryonic fibroblasts (MEFs) isolated from E13.5 mouse embryos mitotically inactivated by Mitomycin C treatment) in 96-well plates in ESC medium + LIF. ESC medium comprises KnockOut-DMEM, 1%FBS (Pan Biotech), 100U/ml Penicillin-Streptomycin (Gibco), 2 mM L-Glutamine), 1x MEM Non-Essential Amino Acids Solution (Gibco) and 0.2 mM 2-Mercaptoethanol (Gibco). Lmx1a-overexpressing clones were grown in puromycin until the start of an experiment. Lmx1a expression was induced using 2µg/ml Doxycycline at t=0h.

A replica plate without feeder cells was used for DNA isolation and genotyping. Knockouts were identified as lines that had large deletions and resulting frameshift mutations in early exons. Absence of gene product was confirmed by immunofluorescence where antibodies were available. Selected clones were expanded on feeder cells in ESC medium for 2-3 passages, then transferred to N2B27+2i+LIF.

### Cell differentiation

For differentiation experiments, ∼2,600 cells per cm^2^ were plated in N2B27 medium + LIF (1,000 U/ml) on CellBIND dishes pre-coated with 0.1% gelatine (Sigma) and incubated overnight. Differentiation was initiated by adding N2B27 medium supplemented with 10ng/ml bFGF for 48 hours, followed by a pulse of 10ng/ml bFGF + 5µM CHIR99021 for 24 h ^25^ . Subsequently, the medium was changed to N2B27 supplemented with 100nM RA and BMP4 at the indicated concentration for the indicated amount of time. Addition of RA + BMP4 is considered as t=0h.

To differentiate cells on stencils, PDMS stencils were fabricated to fit 2-well µ-slide ibiTreat dishes following a custom protocol ^27^ based on ^58^. Prior to plating cells on stencils, 1.5–1.8 million cells were plated on 100 mm CellBIND dishes and cultured overnight in N2B27 medium + LIF (1000 U/ml), and then in N2B27 medium + 10ng/ml bFGF for 24h. Cells were then replated on stencils on the second day of bFGF treatment. For this, stencils were placed into the wells of 2-well µ-slide ibiTreat dishes pre-coated with 0.1% gelatine (Millipore) and dried for 30 min. Stencils were covered with a 1:1 mix of Neurobasal Media (Gibco) and DMEM/F12 (Gibco) and dishes were placed into a desiccator to remove air bubbles. After 24 h in N2B27 + bFGF, cells were dissociated from the 100 mm dishes and 2.8-3M cells per well were seeded on the prepared 2-well dishes in N2B27 medium with 10ng/ml bFGF + 10µM Y-27632 ROCK inhibitor (Tocris) media for 3h. Cells were then washed and further cultured in N2B27 + 10ng/ml bFGF. The next day, the stencils were removed, and the differentiation protocol continued with the media change as described above from day 3 onwards.

To differentiate cells on restricted micropatterned surfaces (Fig. S2D), we used micropatterned glass chips Arena A 500µm diameter (Cytoo). Chips were coated overnight with 1:40 dilution of laminin (Sigma) in PBS in a humid chamber, then placed into a 6-well dish and washed 2x with PBS. Similar to plating on stencils, 3M cells on the second day of bFGF treatment were plated onto the chips in N2B27 medium with 10ng/ml bFGF + 10µM Y-27632 ROCK inhibitor (Tocris) and washed 3h after plating.

To inhibit BMP signalling, medium was supplemented with 1µM LDN-193189 (Tocris) as indicated.

### Immunostaining and imaging

Cultured cells were fixed for 18 min on ice with cold 4% PFA. Samples were incubated in PBST (PBS + 0.1% Triton X-100) 3x 5-10 min, then 2-3h in blocking buffer (PBST + 1% BSA) at room temperature, followed by incubation with primary antibodies overnight at 4°C, 3 washes 5-10min each in PBST, secondary antibodies and DAPI (Sigma) for 2h at room temperature, 3 washes 5-10min each in PBST. Samples were stored in Ibidi mounting medium (Ibidi). Antibodies used are listed in Table T3.

**Table T3.** Antibodies.

| Name | Species | Source | Cat. No. | Dilution |
| --- | --- | --- | --- | --- |
| AP-2alpha | mouse | Santa Cruz | SC-12726 | 1:200 |
| Atoh1 | rabbit | Proteintech | 21215-1-AP | 1:2000 |
| LMX1A | rabbit | Sigma | HPA030088 | 1:2000 |
| MASH1 (ASCL1) | mouse | BD Pharmingen | 556604 | 1:100 |
| Olig3 | mouse | Abcam | ab168573 | 1:200 |
| Phospho-Smad1/5 | rabbit | Cell Signalling Technologies | 9516 | 1:800 |
| Sox2 | goat | R&D | AF2018 | 1:100 |
| Sox2 | mouse | R&D | MAB2018 | 1:500 |
| Sox9 | rabbit | Cell Signalling Technologies | 82630 | 1:150 |
| Sox10 | mouse | Santa Cruz | sc-365692 | 1:50 |
| Wnt1 | rabbit | Abcam | ab15251 | 1:100 |
| ErbB3 | sheep | R&D | AF4518 | 1:500 |
| GFP | sheep | AbD Serotec | 4745-1051 | 1:1000 |
| T/Brachury | goat | R&D | AF2085 | 1:250 |
| Dbx1 | rabbit | <sup>65</sup> |  | 1:10000 |
| Olig2 | goat | R&D | AF2418 | 1:100 |
| Nkx6.1 | mouse | DSHB | F55A10 | 1:50 |

**Table T4.** Kits.

| Name | Source | Cat. No. |
| --- | --- | --- |
| PureLink RNA Mini Kit | Invitrogen | 12183025 |
| PureLink DNase Set | Invitrogen | 12185010 |
| SuperScript III First-Strand Synthesis | Invitrogen | 18080051 |
| Phusion High-Fidelity DNA Polymerase | New England Biolabs | M0530 |
| Q5 High-Fidelity DNA Polymerase | New England Biolabs | M0491 |
| T4 DNA Ligase | Takara | 2011A |
| In-Fusion HD Cloning Kit | Takara | 639649 |

**Table T5.**
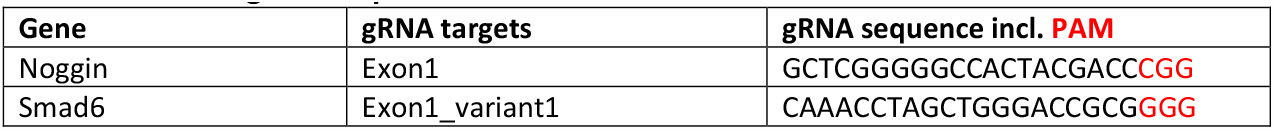
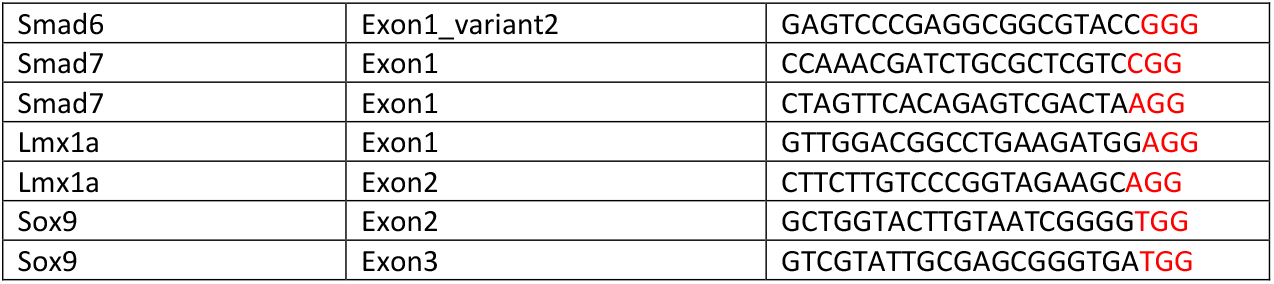
CRISPR gRNA sequences.

Mouse embryos were fixed and cryoprotected as previously described ^59^. For immunostaining of transverse sections, slides were incubated for 20min at 42°C in PBS, then in PBS + 0.1% Tween + 1% BSA for 2h at room temperature, in primary antibody solution overnight at 4°C, washed 3x for 5-10min with PBS + 0.1% Tween, in secondary antibodies and DAPI for 2h at room temperature, then washed 3x for 5-10min with PBST and mounted in ProLong Antifade Mounting medium (Thermo Fisher Scientific).

For whole mount embryo preparations, E8.5 embryos were fixed for 50min in 4% PFA then washed 3x 5-10min in PBS+0.1% Tween, and dehydrated using a Methanol series (25, 50, 75, 100%) and stored at -20°C. For immunostaining, embryos were rehydrated, washed in PBST and incubated in primary antibody at 4°C overnight, followed by 3x 5min washes in PBST and a long wash overnight, followed by overnight incubation in secondary antibody solution, 3x 5-10min washes and another overnight wash. Embryos were mounted in Prolong Antifade Mounting Medium.

Images of ESC colonies were acquired on an inverted Zeiss LSM800 confocal microscope with GaAsP PMTs, using 10x/20x objective, z-stacks with at least 15% overlap. Whole mount HCRs were imaged on a Nikon Spinning Disk CSU-W1 with a pinhole size of 25µm using a CFI Apo LWD λ S 20x WI objective with water. Tile scans were stitched using Imaris stitcher.

### Hybridisation chain reaction (HCR)

HCR probes were ordered from Molecular Instruments. For HCR on 2-well ibidi slides, the “Protocol for mammalian cell on a chambered slide” (https://www.molecularinstruments.com/hcr-rnafish-protocols, Revision Number: 4) was followed with minor modifications. Briefly, cells were washed with PBS, fixed with 4% PFA for 20min on ice, washed 3x with PBS, and stored in 70% EtOH at -20°C overnight to a maximum of 3 weeks. EtOH was aspirated and samples were dried for ∼30min at room temperature. After 2 washes with 2x SSC, samples were prehybridised in probe hybridization buffer at 37°C for 30-60min. Samples were incubated with 1.2pmol of each probe at 37°C overnight. Following 4x 5min washes with probe wash buffer at 37°C and 2x 5min washes with 5xSSCT at room temperature, samples were pre-amplified in amplification buffer for 30-60 min at room temperature. Samples were incubated overnight with 18pmol of each snap-cooled hairpin in amplification buffer at room temperature. After 2x 5min washes with 5x SSCT at room temperature, samples were incubated for 5min in 5x SSCT with DAPI, followed by another 2x 5min washes with 5x SSCT. Ibidi mounting medium was added to the samples that were then stored at 4°C prior to imaging.

For HCR on tissue sections, “In situ HCR v3.0 protocol for sample on slide” (https://www.molecularinstruments.com/hcr-rnafish-protocols, Revision Number: 4) was followed with minor modifications. Briefly, embryos were dissected in cold PBS and fixed depending on developmental stage (E8.5: 50min, E9.5: 55min, E10.5: 70min, E11.5: 90min) in 4% cold PFA on ice. Embryos were washed 3x 5min in cold PBS and incubated in 15% sucrose at 4°C for 1-6h. Brachial regions were further incubated in 30% sucrose (except for e8.5 where whole embryos were incubated) for 2h-overnight. The tissue was then incubated in a 1:1 mix of 30% sucrose:OCT for 1-2h, then in OCT for 1-2h, then transferred to a mould, and frozen on dry ice. OCT blocks were cryosectioned into 14µm thick sections and stored at -80°C. For HCR, slides were air-dried for 5-10min and postfixed for 15min in 4% PFA. Slides were washed 2x 10min in cold PBS and then incubated in 70% EtOH for 10min on ice, followed by incubation in 70% EtOH for 4h - overnight at -20°C. Slides were washed 3x in Hybridisation Wash Buffer (10% Formamide in 2x SSC), dried and then incubated in probe hybridization buffer for 10min at 37°C in a humidified chamber. The tissue was incubated overnight in hybridization buffer containing 0.4pmol of each probe mixture, which was covered with Parafilm to prevent evaporation. Slides were washed in probe wash buffer for 5min, then increasing % of 5x SSCT in probe wash buffer 15 min each until 100% 5x SSCT 15min, all at 37°C, and 5x SSCT for 5min at room temperature. Slides were dried and incubated in amplification buffer for 30min at RT, followed by incubation with 6pmol of each hairpin overnight, covered with Parafilm to prevent evaporation. Following 1x 5min wash with 5x SSCT + DAPI, 2x 30min 5x SSCT, 1x 5min SSCT, all at RT, the tissue on the slide was covered with Prolong Gold and a glass coverslip.

For HCR of whole-mount mouse embryos, “HCR RNA-FISH protocol for whole-mount mouse embryos” (https://www.molecularinstruments.com/hcr-rnafish-protocols, Revision Number: 9) was followed with minor modifications. Briefly, e8.5 mouse embryos were dissected in cold 4% PFA and fixed overnight at 4°C. Embryos were washed 2x 5min with PBST, dehydrated with a series of graded methanol/PBST washes on ice 10min each, then stored in 100% methanol at -20°C. After rehydration to PBST with a series of graded methanol/PBST washes on ice 10min each, embryos were washed 2x 5min in PBST at room temperature and then incubated in 10µg/ml proteinase K in PBST for 10min at room temperature. Embryos were washed 2x 5min with PBST, postfixed with 4% PFA for 20min at room temperature, and again washed 3x 5min with PBST. Embryos were incubated first for 5min, then for 30min in probe hybridization buffer at 37°C, and then incubated overnight in probe hybridization buffer containing 2pmol of each probe set. This was followed by 4x 15min washes with probe wash buffer at 37°C, 2x 5min washes with 5x SSCT at room temperature, 1x 5-30min in amplification buffer at room temperature, incubation overnight in amplification buffer containing 30pmol of each hairpin. On the next day, embryos were washed 2x 5min in 5x SSCT, 1x 30min in 5x SSCT+DAPI, 1x 30min in 5x SSCT, 1x 5min in 5x SSCT and then mounted on a glass slide.

### qRT-PCR

RNA was extracted from cells cultured in 12-well plates using the Purelink RNA Mini Kit (Invitrogen). DNA was removed by incubating the preparation on the column with Purelink DNase (Invitrogen). RNA was quantified and stored at -80°C. cDNA was generated from 500-2000ng total RNA using SuperScript III (Invitrogen) with random hexamer primers. Quantitative RT-PCR was performed in triplicates using Roche Lightcycler 480 SYBR Green in 384-well format.

### RNA sequencing sample preparation

Cells differentiated on stencils were dissociated into single cells and pelleted by centrifugation. Three biological replicates per timepoint and condition were collected. Sequencing libraries were prepared by the VBC Sequencing facility using NEB poly-A stranded kit. Pooled libraries were sequenced on the NextSeq2000 P3 (paired-end 50bp reads).

### RNA sequencing data analysis

RNA sequencing reads were trimmed using Trim galore v0.5.0 (https://github.com/FelixKrueger/TrimGalore), aligned to the GRCm38 genome using STAR aligner v2.6.0c (https://github.com/alexdobin/STAR) and counted using GeneCounts. Gene count tables were imported into R (v4.3.0) (R Core Team) and converted to a DESeq data set using DESeqDataSetFromMatrix (https://bioconductor.org/packages/release/bioc/html/DESeq2.html). Log2 fold change relative to BMP4 concentration 0 ng/ml or time =0h (as indicated) was calculated using DESeq2 ^60^ see library in R. Heatmaps were generated using the pheatmap library in R. RNA-sequencing data generated in this study is available at GEO (Gene Expression Omnibus, https://www.ncbi.nlm.nih.gov/geo/), accession number GSE247069.

### Mouse strains

All work with animals was approved under the license BMWFW-66.018/0006-WF/V/3b/2016 from the Austrian Bundesministerium für Wissenschaft, Forschung und Wirtschaft. All procedures were performed in accordance with the relevant regulations. CD-1 outbred mouse strain (Charles River) was used to obtain wildtype embryos. Transgenic strains were maintained onto CD-1 background. The following strains and respective genotyping protocols were previously described: Wnt1-Cre2 (JAX 022501, ^61^), Sox2CreERT2 (JAX 017593, ^62^), Nog^Flox^ (JAX 016117, ^63^). Pregnant mothers were intraperitoneally injected with 4mg tamoxifen in sunflower oil at the indicated stages. Embryos were dissected and fixed according to developmental stage, and embedded in gelatine for cryosectioning. Yolk sacs were collected for genotyping.

### Quantification of fluorescence intensity profiles in differentiated cells

Fluorescence intensity (FI) profiles in maximum intensity projection images of antibody or HCR stainings of differentiated colonies were measured as a function of the radial distance from the colony centre using a custom script in Python 3 (https://github.com/dbrueckner/NeuralTubeColonies). Specifically, we identify the position of the colony centre ***X***_*c*_ using the Sox2 FI by taking the mean position ***X***_***i***_ = (*x, y*) of all pixels multiplied by their respective Sox2 FI *FI*_Sox2_(***X***_*i*_): i.e.

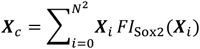

for an image with N x N pixels with indices *i*. Thus, the radial distance of every pixel from the inferred colony center is 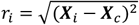. The radial fluorescence intensity profile of the protein or gene of interest is then measured using the average FI as a function of the radial coordinate *f*_α_(*r*) =⟨*FI*_α_(***X***)|*r* = *r*_*i*_⟩. Colonies that were strongly asymmetric due to distortions during the culture or sample preparation process were excluded from analysis.

After the spatial FI profiles were measured, the maximum intensity value of each profile was identified and used as a data point in further analysis. Within a given experiment that includes different conditions or time points, all colonies were processed and imaged in an identical manner and therefore the fluorescence intensities are comparable. Within each experiment, background FI was determined from a time point or condition where there was no expression of the analysed marker, and subtracted from data. To combine data from independent experiment repeats, data points were normalized to the experiment mean for a defined time point (in most cases we used the time point of maximal FI across the timecourse) or condition (in most cases, we used 0.5ng/ml BMP4).

To obtain an estimate of the number of cells expressing a given protein of interest, we measured the area occupied by cells positively stained for that protein in maximum projection images. Threshold fluorescence intensities were determined using a negative control sample with a custom script in Python 3 using Matplotlib’s imshow() and applied to all images of the same experiment.

### Quantification of fluorescence intensity profiles in embryos

HCR wholemount images were analysed in Fiji ^64^. Measurements were obtained by tracing the edge of the Sox2+ neural plate in maximum intensity projections using the freehand line tool with width set to 12µm. FI profiles were obtained using the plot profile function. Profiles were smoothed using a rolling mean with a window size of 10. Profiles were normalised to the maximum value of each trace and background subtracted. For Lmx1a and Bra, background was defined as the minimum value of each profile. For pSmad1/5 and Nog, the background was estimated using regions of the embryo that did not express these markers.

To quantify RP size from images of immunostained sections, images were first cropped to exclude Sox2 cells outside of the neural tube and Lmx1a cells outside the dorsal midline region. Lmx1a expressing region at the dorsal midline and the Sox2 positive region of the neural tube were then determined by automated processing using Python 3. For this, a FI threshold was determined as the mean (for Lmx1a) or mean – 1*SD (for Sox2) of FI intensity values above 10% of the maximum FI. Saturated pixels were excluded from the analysis. The expression area was considered the area above the threshold. Features smaller than 8 pixels were considered noise and removed. The Lmx1a positive area was then normalised to the Sox2+ positive area.

## Acknowledgements

We thank J. Briscoe for comments on the manuscript. Work in the AK lab is supported by ISTA, the European Research Council under Horizon Europe: grant 101044579, and Austrian Science Fund (FWF): F78 (Stem Cell Modulation). SR is supported by Gesellschaft für Forschungsförderung Niederösterreich m.b.H. fellowship SC19-011. D.B.B. was supported by the NOMIS foundation as a NOMIS Fellow and by an EMBO Postdoctoral Fellowship (ALTF 343-2022).

## Supplementary Information

### Reaction-diffusion model of BMP signaling in 2D stem cell colonies

#### 1 Reaction-diffusion model of BMP signaling

To capture BMP signaling dynamics in a minimal biophysical model, we constructed step-by-step the simplest reaction-diffusion network that is consistent with experimental observations. Making use of the radial symmetry of the system, we write a reaction-diffusion system of equations only as a function of time and of the radial coordinate *r*. We simulate the system on a large domain 0 < *r* < *R*_∞_. To implement the cell colony, we restrict all production terms to only be non-zero within the colony of radius *R* (by multiplying the production terms by the Heaviside function Θ(*R*−*r*)). In contrast to production, diffusion and degradation can also take place outside the colony. As discussed in the main text, our model considers the dynamics of pSmad1/5 activity (here abbreviated as pSmad), the diffusible ligand (BMP), BMP inhibitor (BMPi) as well as the transcription factor Lmx1a. Note that while BMP and BMPi are diffusible (with diffusion coefficients *D*_*b*_ and *D*_*i*_), the intracellular species (pSmad, Lmx1a) cannot diffuse.

We implement the hypothesized interaction network for {BMP, BMPi, pSmad, Lmx1a}, whose concentrations are denoted as {*b*(*r, t*), *n*(*r, t*), *s*(*r, t*), *l*(*r, t*)}, as a set of partial differential equations. Before we discuss the specific implementation of the reaction network in the following sections, we first define the general set of equations as well as initial and boundary conditions that are used throughout. Specifically, we use the following equations:

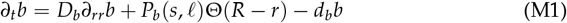

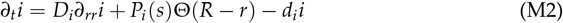

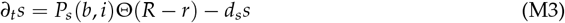

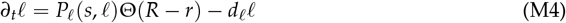

where *P*_*k*_ and *d*_*k*_ are the production functions and degradation rates of species *k*. The dependencies of the production functions *P*_*k*_ are motivated by the known properties of the species (see main text). Specifically, pSmad is activated by BMP and BMPi (such as Noggin) inhibits the BMP-mediated pSmad activation. pSmad activates the production of both BMP and BMPi. In addition, pSmad activates the production of Lmx1a, and Lmx1a activates BMP. Throughout, we solve this model subject to the following boundary conditions

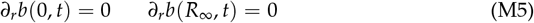

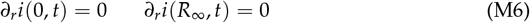

using *R*_∞_/*R* = 10 for computational simplicity. Throughout, we non-dimensionalize the system to minimize the number of parameters. Thus, we remove the spatial units by setting *R* = 1, so that all spatial scales (i.e. in the diffusion coefficients) are measured in units of the colony size. In contrast, we retain the units of time in order to make predictions of the temporal dynamics in units of hours, which can then be directly compared to experiments. Therefore, the simulations are initialized at *t* = 0 and run up to a final time *t* = 96 h as in experiments. Since the system is subjected to a spatially homogeneous exogeneous BMP input at *t* = 0 with varying concentration denoted as *b*_0_, we use the following initial condition for BMP:

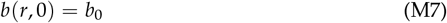

In experiments, there is no expression of pSmad or Lmx1a at *t* = 0 (Fig. 1D,F and Fig. 2A,C). We therefore use the initial conditions:

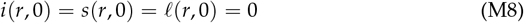

Together, Eqs. (M1) – (M8) fully specify the reaction-diffusion system.

#### 2 First phase dynamics

To gain insight into how the initial up- and down-regulation of pSmad is regulated, we first study the three-component network for {BMP, BMPi, pSmad}, which provides a model for the first phase of patterning dynamics (Fig. M2A). To reduce the number of parameters, we first explore the simplest implementation with linear reaction kinetics for BMP and BMPi, and a generic Hill function for pSmad activation arising from a balance of BMP and BMPi levels:

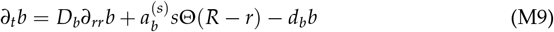

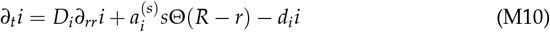

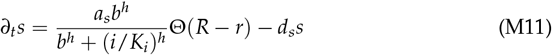

where 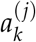 is the activation rate of species *k* by species *j, K*_*i*_ is the threshold at which pSmad production becomes sensitive to BMPi concentration, and we use *h* = 2 throughout. To reduce the number of parameters in the model, we remove the dimensions of each species’ concentration, since we are interested in comparing the behaviour of normalized concentrations to the normalized concentrations in the experiment.

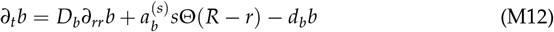

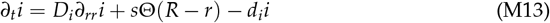

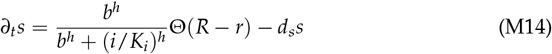

Solving these equations yields concentration profiles that peak at the colony center, as BMP and BMPi are produced only within the colony, but are free to diffuse and degrade outwards. This is in contrast to the experimentally observed spatial profiles, where pSmad peaks at the colony boundary. Previous work has suggested that in 2D cell colonies cultured on micropatterned surfaces, cells at the boundary have higher sensitivity to BMP ligands [1]. To incorporate this, we postulate that the activation of pSmad by BMP is stronger near the colony edge. We therefore modify Eq. (M14) as follows:

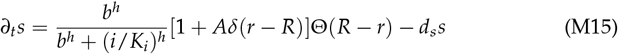

where we take *δ*(*x*) = exp[−*x*^2^/(2*σ*^2^)]. We find that using *A* = 1 and *σ* = 0.1, we recover pSmad profiles qualitatively similar to the experiment (Fig. M1).

**Figure M1:**
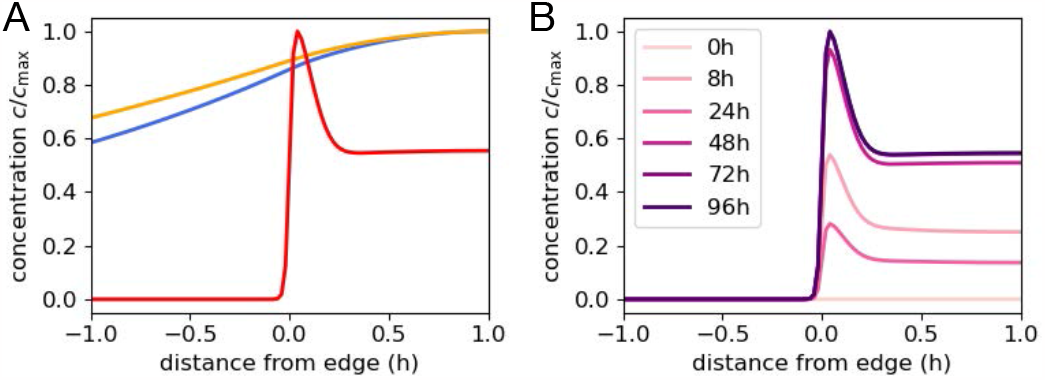
Spatial profiles. (A) Profiles showing BMP, BMPi and pSmad at the final time-point (*t* = 96h). (B) Profiles of pSmad over time. Both plots correspond to the full dynamics including Lmx1a (section 3.3).

##### 2.1 Steady-state analysis of the first phase network

Based on the simulations of Eqs. (M12)-(M14), we find that the spatial profile of pSmad has a near-constant length scale over time, consistent with experiments. Thus, the temporal dynamics of the circuit are captured by the maximum levels of pSmad signaling at every time point. These dynamics are approximately described by a set of ordinary differential equations for the concentrations at the colony edge, which allows us to solve analytically for the steady-state concentrations as a function of the different model parameters:

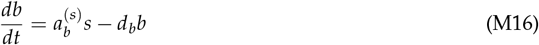

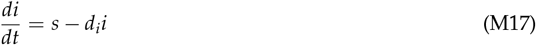

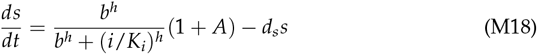

In the experiment, the system is subjected to an initial exogeneous BMP concentration to which it responds by pSmad activation and subsequent production of endogeneous BMP and BMPi. Within the framework of our model, we are therefore looking to characterize the temporal response of the system to the initial condition Eq. (M7) and its subsequent approach to a steady state denoted by *b*^*\**^.

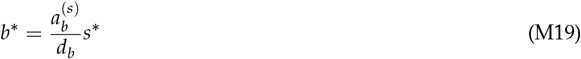

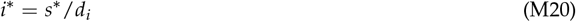

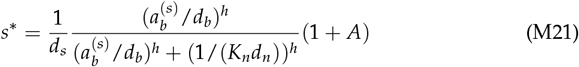

We plot the dependence of the steady-state concentrations as a function of parameters (Fig. M2B-D), which show that there are three possible parameter regimes with qualitatively different behaviours:

- Regime I: zero-steady state: *b*^*\**^ *≈* 0. For very low BMP production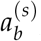, the steady-state BMP concentration is nearly zero in this model.
- Regime II: non-zero steady state approached from below: *b*_0_ < *b*^*\**^. At larger values of BMP production 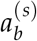, the BMP steady state is non-zero and can be approached from below, i.e. the initial exogeneous BMP concentration is lower than its steady-state value.
- Regime III: non-zero steady state approached from above: *b*_0_ *> b*^*\**^ *>* 0. Alternatively, in the non-zero steady state regime, the initial condition may be larger than the steady state BMP concentration.

**Figure M2:**
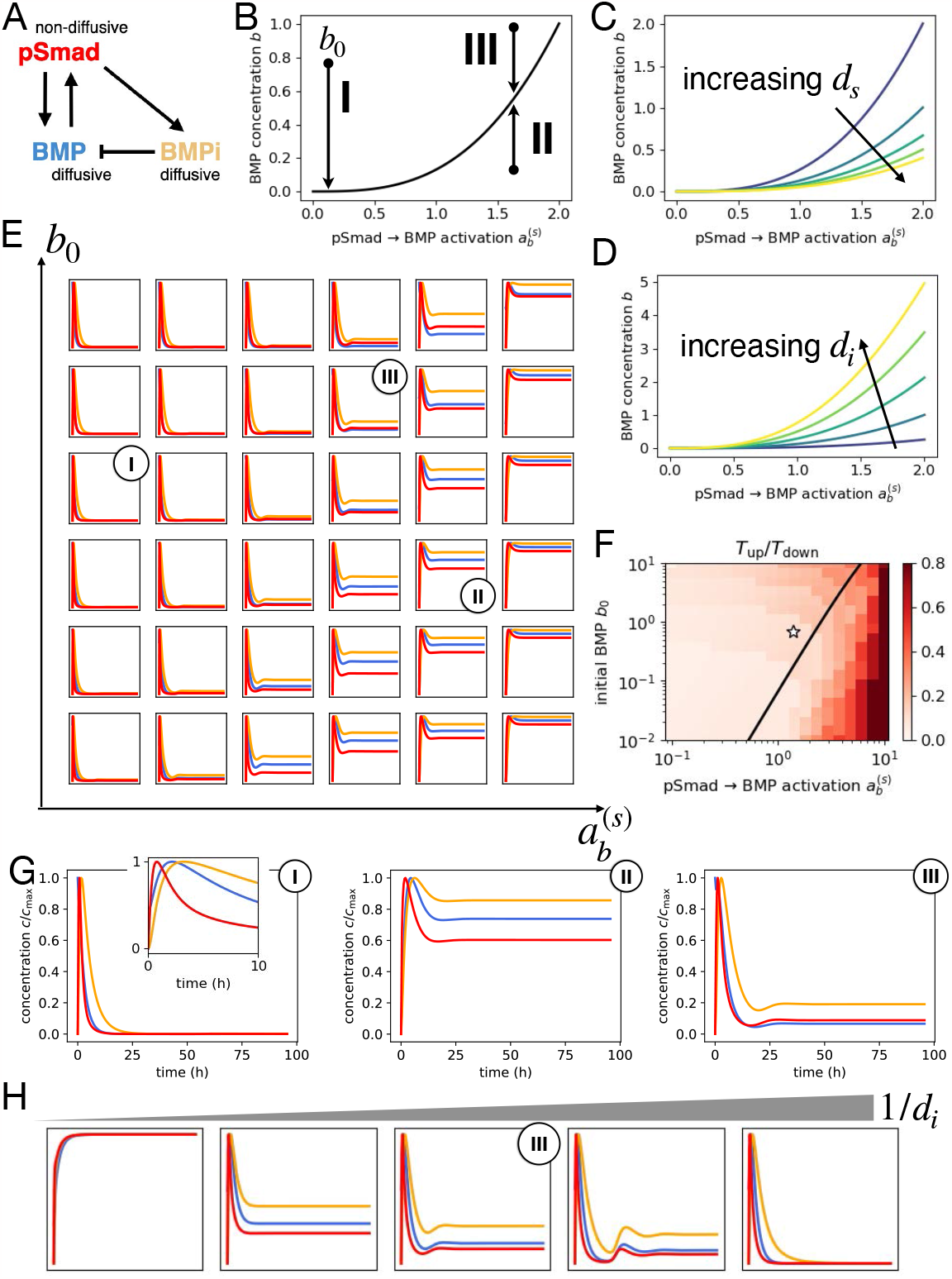
Dynamics of the first phase network. (A) Schematic of the first phase network. (B) Steady-state BMP concentration as a function of the pSmad to BMP activation rate 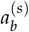, with the three parameter regimes indicated, obtained by combining Eqs. (M19) and (M21). (C,D) Steady-state BMP concentration as a function of the pSmad to BMP activation rate 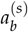 for varying *d*_*s*_ and *d*_*i*_, respectively. (E) Dynamics of the first phase network as a function of the parameter space 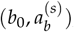), showing rapid upregulation of pSmad signaling in all regimes, but no second phase peak. (F) Ratio of time to maximum pSmad *T*_up_ to time to downregulation to subsequent minimum *T*_down_ as a function of Parameters 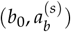. (G) Detailed plots corresponding to parameter regimes I, II and III. (H) Dynamics of the first phase network as a function the inhibitor time-scale.

Based on simulations for the full spatio-temporal model (Eqs. (M12), (M13), (M15)), we find that the steady-states of this model are in agreement with the simplified analysis above (Fig. M2E). Importantly, we find that the timing of the pSmad maximum is robust across these three regimes, as it is determined by the relative delay between BMP and BMPi concentrations (Fig. M2E, inset Fig. M2G).

However, the three parameter regimes exhibit quantitatively different downregulation behaviour from the first phase peak. In regime I, the peak rapidly decays to a vanishing pSmad concentration (Fig. M2G, I). In regime II, the initial peak is only weakly downregulated as the initial overshoot is close to the final steady state (Fig. M2G, II). In regime III, a rapid pSmad upregulation is followed by strong downregulation to a small but finite steady-state concentration (Fig. M2G, III). These observations are quantified in Fig. 4C and Fig. M2F.

Our model of the first phase network should capture three key experimental observations: (i) the first phase peak exhibits rapid upregulation with strong subsequent downregulation. (ii) a small but finite pSmad concentration is observed at long times in the Lmx1a knockout experiment (which corresponds to the first phase network in the model). (iii) We do not observe endogeneous BMP production in the system at early time-points (Fig. S3C).

Based on the time-scale parameters of BMP and BMPi signaling (i.e. their degradation rates), observation (i) is captured in all three regimes as the initial BMP activates BMPi, leading to a rapid downregulation of pSmad. Observation (ii) implies that the system is in regime II or III, corresponding to a non-zero steady state. Finally, observation (iii) suggests that the system is in regime III, where the BMP amplitude decays to steadystate levels, rather than initially increasing due to endogeneous production. Thus, we use parameters corresponding to regime III.

Crucially, none of the three parameter regimes can result in oscillations in which the second peak has a similar amplitude as the first one, as observed in the experimental dynamics, even when additional parameters such as the inhibitor time-scale are varied (Fig. M2H).

#### 3 Second phase network: including Lmx1a dynamics

##### 3.1 Lmx1a dynamics is non-linear

Since a minimum network including BMP, BMPi and pSmad could capture the first, but not the second phase of the experimental dynamics, we explored the possibility that Lmx1a plays a key role in mediating the upregulation of pSmad in the second phase. As discussed in the main text, we incorporate the well established observation that Lmx1a activates BMP ligand production [2,3] (Eq. (M22)). Furthermore, we incorporate the observation that BMP signaling can promote Lmx1a expression [4] (Eq. (M30)). To again start with the simplest approach, we first explore a linear dependence of Lmx1a production on pSmad levels:

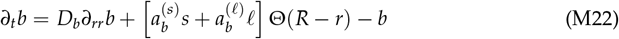

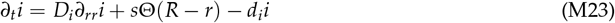

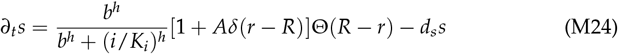

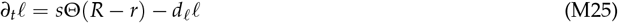

This introduces two additional free fitting parameters: the degradation rate of Lmx1a *d*_*l*_, which determines the time-scale of its response; and the activation rate of BMP by Lmx1a 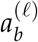. We therefore vary these parameters to test whether this model can capture the experimental dynamics.

**Figure M3:**
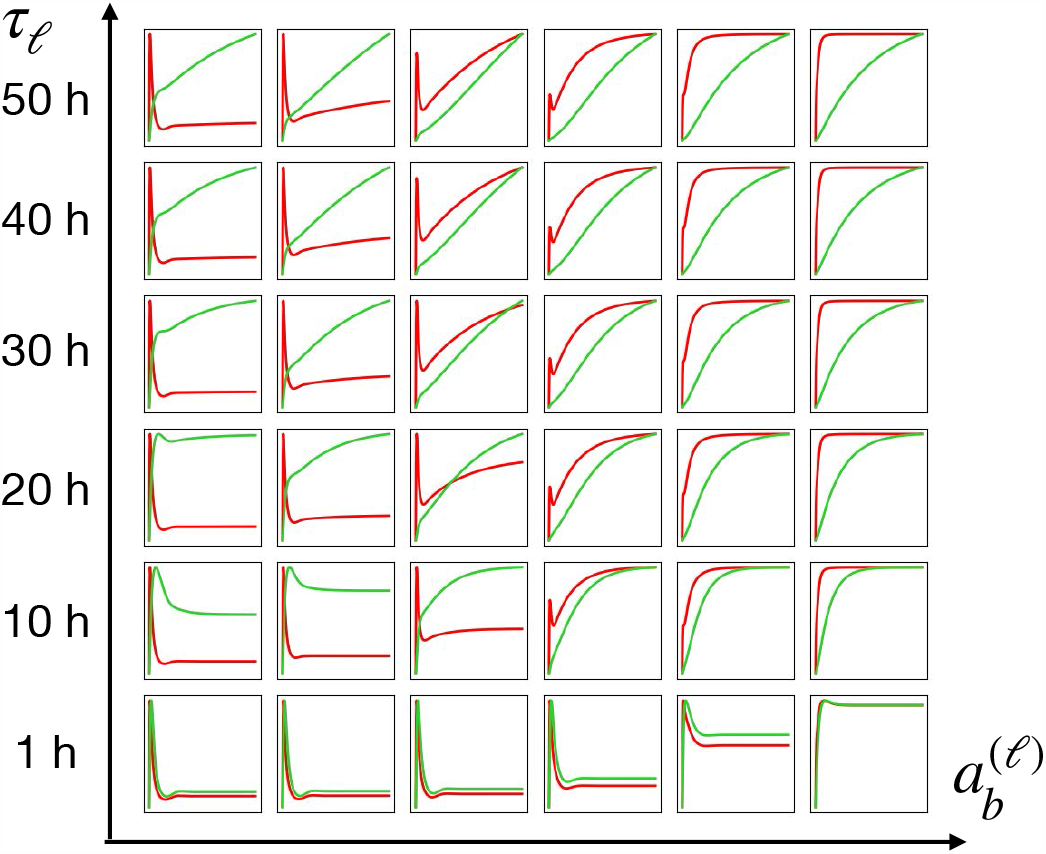
Dynamics of linear Lmx1a activation. Dynamics of pSmad (red) and Lmx1a (green) as a function of Lmx1a parameters 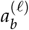 and *τ*_*l*_ = 1/*d*_*l*_.

Interestingly, at large Lmx1a time-scales *τ*_*l*_ = 1/*d*_*l*_ ≳ 20h, we find that this model captures the basic qualitative separation of the two phases of signaling, with a first pSmad peak, and a second phase with slowly increasing pSmad levels (Fig. M3). However, this model fails to quantitatively capture other features of the data: Lmx1a concentrations are predicted to rise monotonically from *t* = 0, whereas in experiments, Lmx1a protein is not observed before 24h (Fig. 1D, F). This is a fundamental limitation of this linear model: given that the maximal intensity of the pSmad levels are similar in the first and second phases, a linear relation between pSmad and Lmx1a would always predict similar production functions. Therefore, we next investigate a potential nonlinearity in the Lmx1a dynamics to better capture the experimental observations. To this end, we first directly infer the key time-scale parameter *τ*_*l*_ from experiments to constrain our model search.

##### 3.2 Inference of time-scale parameters from experimental data

In order to measure *τ*_*l*_, we experimentally inhibited BMP signaling using LDN193189 inhibitor at *T* = 72h, and monitored the subsequent decay of pSmad and Lmx1a levels. Since LDN disrupts all BMP signaling, this is equivalent to setting *a*_*s*_ to zero at *T* = 72h in our model and monitoring the subsequent decay dynamics. These are then approximately given by

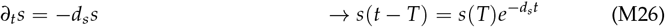

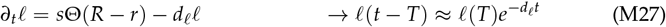

where the Lmx1a decay is exponential if the remaining pSmad rapidly decays without further activating Lmx1a (i.e. if *d*_*s*_ ≫ *d*_*l*_). Fitting these time-scales, we find a strong separation of time scales, with pSmad levels decaying to background levels within an hour, while Lmx1a decays with a time scale of around 10h (based on exponential fits to Fig. 4H):

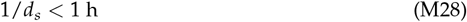

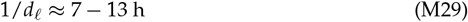

##### 3.3 Nonlinear Lmx1a dynamics

Given the inability of the linear model to accurately predict Lmx1a dynamics, we hypothesized that pSmad alone only provides a weak activation of Lmx1a with rate 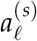, which, beyond a nonlinear threshold *K*_*l*_, self-activates to promote its own production.

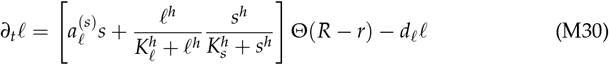

Here, we included a second Hill function dependent on pSmad, thus positing that the Lmx1a self-activation is pSmad dependent. This is required to capture the decay of Lmx1a upon LDN treatment, as it would otherwise continue to self-activate without decay.

This model is the simplest implementation of non-linear dynamics which agrees with the LDN experiment, and adds two key parameters, 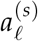 and *K*_*l*_. The time at which the second phase begins depends on the balance of these parameters. Varying both of these parameters, we can closely recapitulate the experimentally observed dynamics in parameter regions where the weak activation of Lmx1a by pSmad crosses the threshold once the first phase peak has decayed (Fig. M4). This cue from pSmad into Lmx1a therefore ensures that the signal is relayed from the first to the second phase.

**Figure M4:**
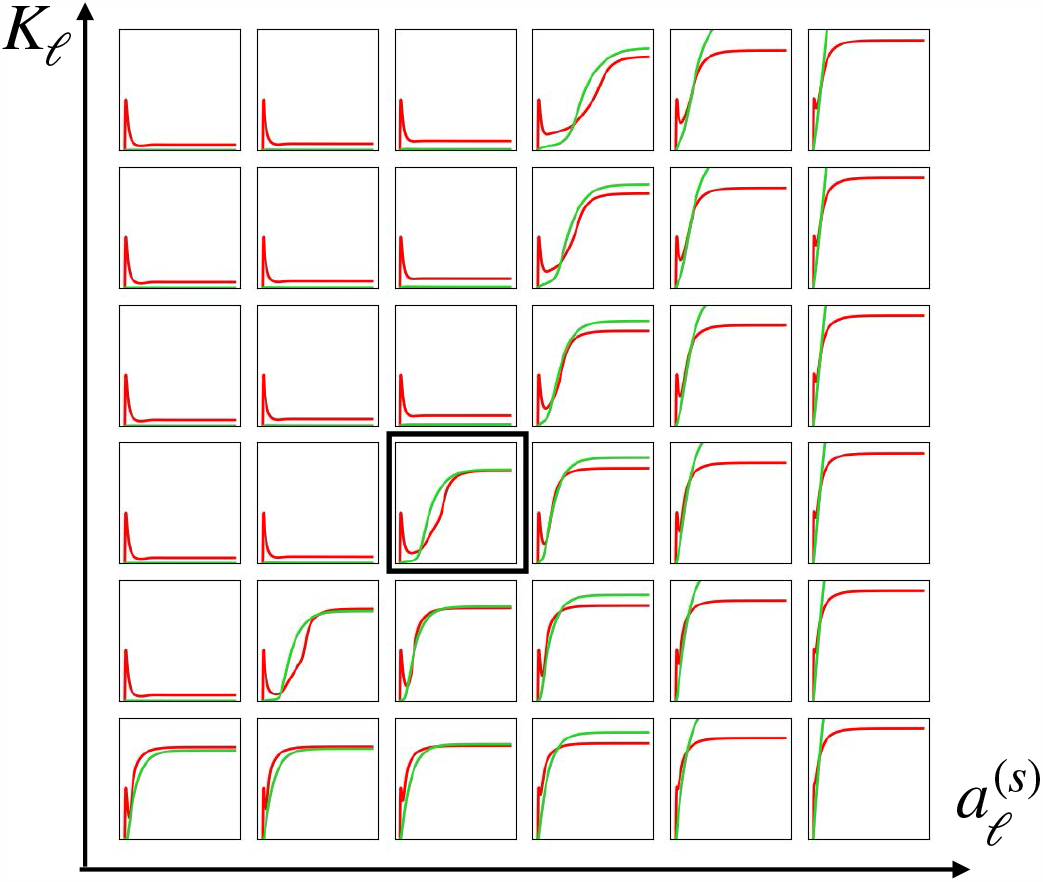
Parameter sweep of nonlinear Lmx1a parameters. Dynamics of pSmad (red) and Lmx1a (green) as a function of Lmx1a parameters 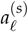 and *K*_*l*_ = 1/*d*_*l*_. Note here all curves are normalized to the same maximum to better show relative changes between parameters.

##### 3.4 Parameter overview

The simulations in sections 2.1, 3.2, and 3.3 constrain the parameters of our model, which are summarized in table M1.

**Table M1:** Overview of parameters used, in units with non-dimensionalized space and concentration coordinates.

| Name | Symbol | Value | Units |
| --- | --- | --- | --- |
| BMP diffusion coefficient | $D_b$ | 1 | $\text{h}^{-1}$ |
| BMPi diffusion coefficient | $D_i$ | 1 | $\text{h}^{-1}$ |
| BMP degradation time-scale | $1/d_b$ | 5 | h |
| BMPi degradation time-scale | $1/d_i$ | 10 | h |
| pSmad degradation time-scale | $1/d_s$ | 1 | h |
| Lmx1a degradation time-scale | $1/d_\ell$ | 8 | h |
| BMP activation rate by pSmad | $a_b^{(s)}$ | 1.4 | – |
| BMP activation rate by Lmx1a | $a_b^{(\ell)}$ | 1 | – |
| Lmx1a activation rate by pSmad | $a_\ell^{(s)}$ | 0.08 | – |
| initial BMP concentration | $b_0$ | 0.7 | – |
| Hill threshold inhibitor | $K_i$ | 0.2 | – |
| Hill threshold Lmx1a self-activation | $K_\ell$ | 1 | – |
| Hill threshold pSmad self-activation switch | $K_s$ | 0.1 | – |

##### 3.5 Predictions of the model

###### 3.5.1 Varying exogeneous BMP concentrations

A key prediction of our model is its behaviour as a function of varying exogeneous BMP (exBMP) concentration. To model the varying initial exBMP concentrations in the experiment, *{*0.5, 1.5, 3, 6*}* ng/ml, we use initial conditions

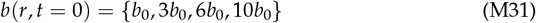

thus using the same relative increase as in the experiment. This predicts that the amplitudes of pSmad peak and minimum levels in the first phase increase with exBMP concentration (Fig. M5A-C). This increase exhibits a saturating trend well predicted by our model due to the nonlinearity in the pSmad inhibition (Eq. (M15)), which implies a maximum rate of pSmad production. The peak amplitude of key BMP inhibitors, including Noggin, exhibit a similar increasing trend with exBMP, as observed experimentally in qPCR (Fig. S4A). In contrast, the model predicts that the duration of the first phase – defined as the time elapsed between the first maximum and the minimum of pSmad concentration – is nearly constant as the exBMP amplitude is increased tenfold (Fig.M5D-F). Specifically, the duration decreases by only *≈* 30%, as compared to an increase in amplitude by *≈* 160%.

**Figure M5:**
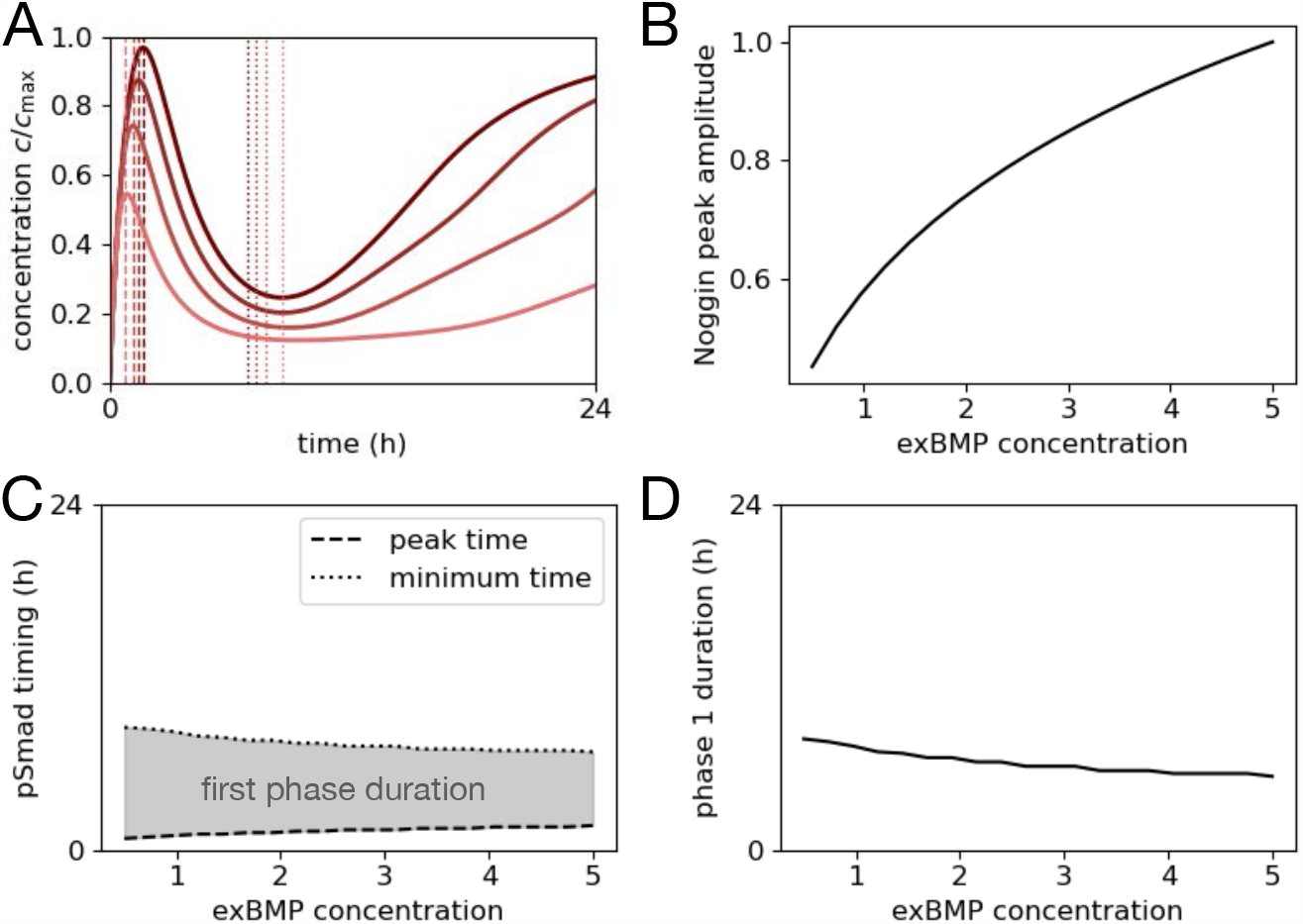
Model predictions for varying BMP concentration. (A) High time-resolution plot of the pSmad dynamics for varying exBMP concentrations. Dashed lines indicate the location of the first phase maximum of pSmad signaling; dotted lines the location of the subsequent minimum. (B) Maximum amplitude of BMPi in the first phase, showing that the model predicts increasing BMPi production with increasing exBMP. (C) Timing of the pSmad peak (dashed) and pSmad minimum (dotted), with first phase indicated in grey. (D) Duration of the first phase, indicating durations relatively insensitive to the exBMP concentration.

###### 3.5.2 BMP inhibitor knockouts

To test the role of the BMP inhibitor species, we simulate a condition in which the production of the BMP inhibitor is reduced by 30%. Specifically, we set 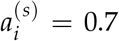 (as defined in Eq. (M10)), compared to its standard value in the non-dimensionalized set of equations of 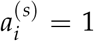. Because we find that several inhibitors of BMP are expressed upon BMP treatment in cultured cells (including Noggin, Smad6/7, Fig. S4), this is a simple model to predict how knocking out one of these inhibitors impacts the dynamics of the system.

Consistent with the model prediction that the second phase of pSmad dynamics critically depends on the Lmx1a subnetwork, we find that the later the inhibitor is removed, the less pronounced its effect on pSmad (Fig. M6). Furthermore, the effect on Lmx1a also strongly declines with time (Fig. S6C).

**Figure M6:**
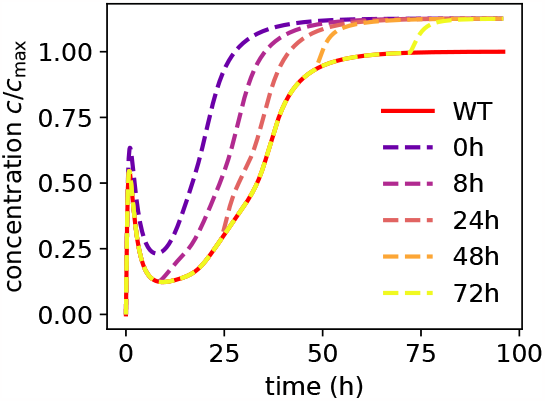
Model predictions for perturbations of the BMP inhibitor at different timepoints. Wildtype dynamics (red line) are compared to reduction of BMPi production rates at different timepoints as shown in the legend (dashed lines).

###### 3.5.3 Relay behaviours predicted by the model

To explore the possible behaviours predicted by our model, we generate phase diagrams of the typical pSmad and Lmx1a relay behaviours as a function of the key parameter combinations.

Specifically, we first investigate behaviours as a function of BMP and Lmx1a activation rates by pSmad, 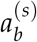 and 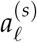, respectively (Fig. M7A). To quantify these behaviours, we use two key aspects of the dynamics. To measure the downregulation of pSmad after the first phase, we define the fractional downregulation *D* = 1−*s*_min_/*s*_max_, where *s*_max_ is the maximum pSmad amplitude in the first phase, and *s*_min_ is the amplitude of the subsequent minimum (Fig. M7B). We use this definition to quantify the degree of downregulation on a unique scale from 0 to 1. Since *s*_min_ < *s*_max_ by definition, and thus 0 < *s*_min_/*s*_max_ < 1, and therefore *D* is between 0 and 1, with *D* = 1 if *s*_min_ = 0 (complete downregulation) and *D* = 1 if *s*_min_ = *s*_max_ (no downregulation).

To measure the produced amount of Lmx1a, we record the Lmx1a concentration at the final time-point, *ℓ*_final_ = *l*(*t* = 96) (Fig. M7C). Based on these two quantities, we observe three typical behaviours (Fig. M7A-C):

1. **first phase only:** if the production rates are too low, the system fails to activate the Lmx1a self-activation loop, leading to failure of the relay mechanism (bottom left corner). In the phase diagram Fig. 4I, we define this phase wherever *ℓ*_final_ *≈* 0.
2. **simultaneous phases:** if the production rates are too high, Lmx1a self-activates before pSmad was downregulated significantly, leading to no clear separation between first and second phase (top right corner). In the phase diagram Fig. 4I, we define this phase wherever pSmad is downregulated by less than 50% of its maximum amplitude, *D* < 0.5.
3. **temporal relay:** between these two phases, we observe a broad parameter regime where a temporal relay with clear ordering of phases, significant pSmad downregulation, and subsequent activation of Lmx1a is observed.

We similarly investigate the behaviours as a function of the BMP activation rates by pS-mad, 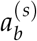, and the initial exogeneous BMP concentration *b*_0_ (Fig. M8A). We find similar qualitative behaviours as in Fig. M7. Star symbols indicate the BMP concentrations in our model for varying exogeneous BMP concentration, which are in a regime where the temporal relay behaviour is robust to changes in the exogeneous BMP concentration. Importantly, all parameter scans are performed in log-scale, meaning that each parameter regime is robust for a broad range of parameters.

**Figure M7:**
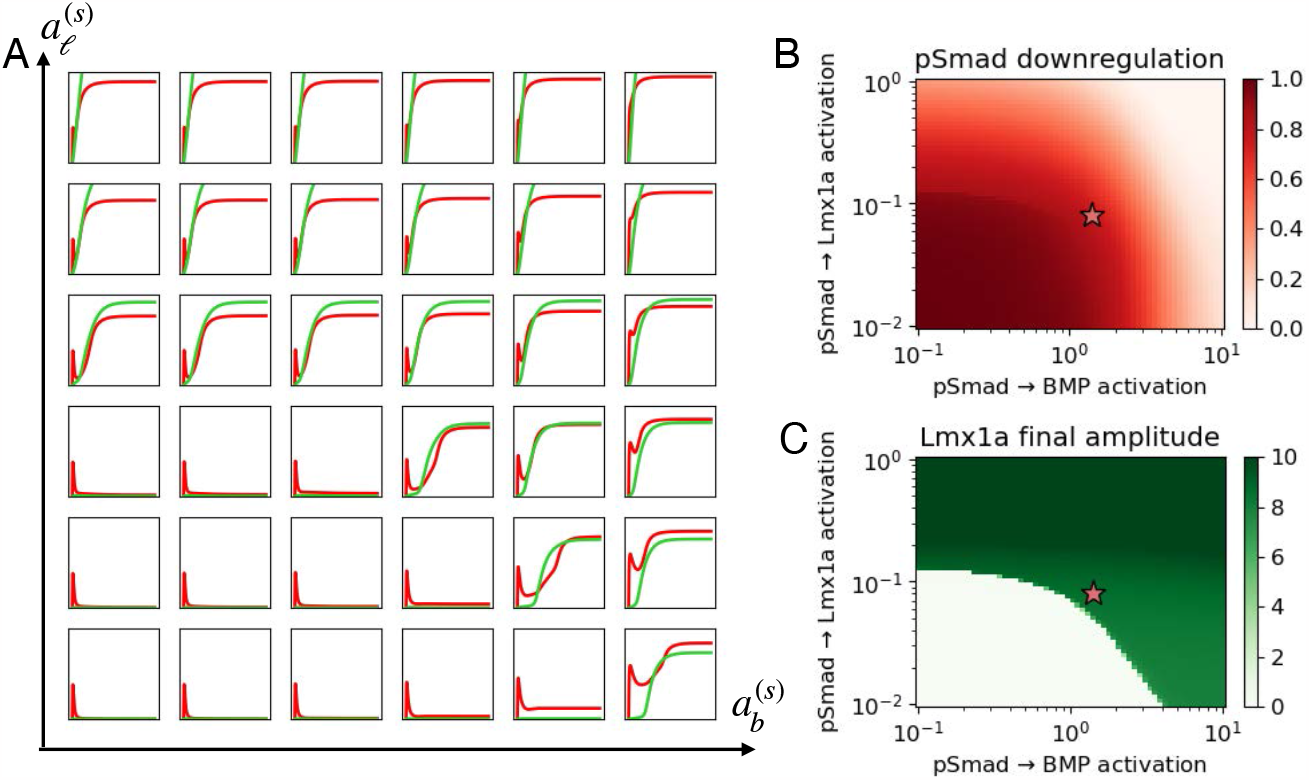
Model predictions for relay behaviours as a function of BMP and Lmx1a production rates. (A) pSmad (red) and Lmx1a (green) concentrations over time as a function of BMP and Lmx1a activation rates by pSmad, 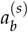 and 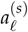. (B) Fractional downregulation of pSmad *D* = 1−*s*_min_/*s*_max_, where *s*_max_ is the maximum pSmad amplitude in the first phase, and *s*_min_ is the amplitude of the subsequent minimum. Star symbol indicates the standard model parameters. (C) Lmx1a concentration at the final time-point, *ℓ*_final_ = *ℓ*(*t* = 96).

**Figure M8:**
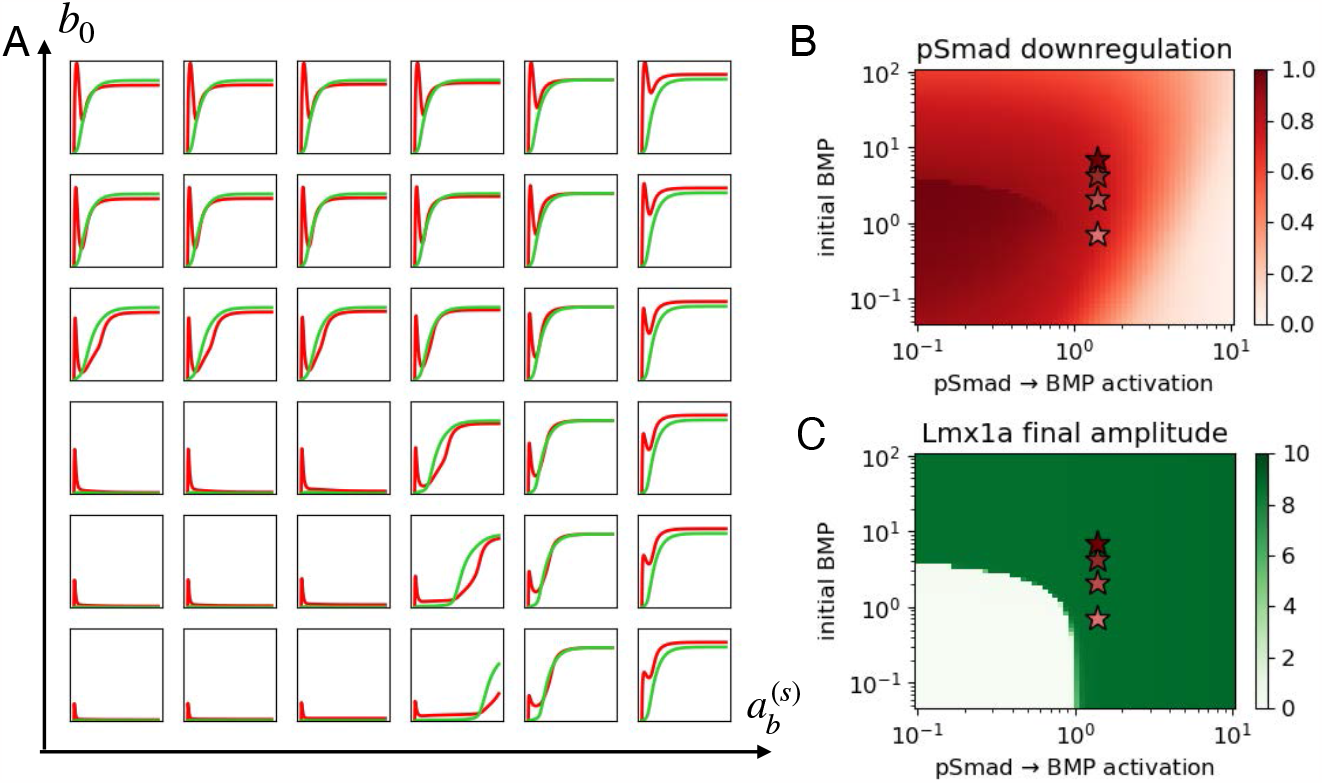
Model predictions for relay behaviours as a function of BMP production rate and initial exogeneous BMP. (A) pSmad (red) and Lmx1a (green) concentrations over time as a function of the BMP activation rates by pSmad, 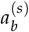, and the initial exogeneous BMP concentration *b*_0_. (B) Fractional downregulation of pSmad *D* = 1−*s*_min_/*s*_max_, where *s*_max_ is the maximum pSmad amplitude in the first phase, and *s*_min_ is the amplitude of the subsequent minimum. Star symbols indicates the standard model parameters for changing exBMP concentrations. (C) Lmx1a concentration at the final time-point, *ℓ*_final_ = *ℓ*(*t* = 96).

